# The sodium leak channel NALCN is regulated by neuronal SNARE complex proteins

**DOI:** 10.1101/2024.09.03.610923

**Authors:** Samuel Usher, Estelle Toulmé, Roberta Florea, Stanislau Yatskevich, Christine C. Jao, Janne M. Colding, Prajakta Joshi, Inna Zilberleyb, Thorsten Trimbuch, Bettina Brokowski, Alexander S. Hauser, Alexander Leitner, Christian Rosenmund, Marc Kschonsak, Stephan A. Pless

## Abstract

The sodium leak channel NALCN is vital for the regulation of electrical activity in neurons and other excitable cells, and mutations in the channel or its auxiliary proteins lead to severe neurodevelopmental disorders. Here we show that the neuronal SNARE complex proteins syntaxin and SNAP25, which enable synaptic transmission in the nervous system, inhibit the activity of the NALCN channel complex in both heterologous systems and primary neurons. The existence of this interaction suggests that the neurotransmitter release machinery can regulate electrical signalling directly, and therefore modulate the threshold for its own activity. We further find that reduction of NALCN currents is sufficient to promote cell survival in syntaxin-depleted cells. This suggests that disinhibited NALCN may cause the puzzling phenomenon of rapid neuronal cell death in the absence of syntaxin. This interaction may offer opportunities for future drug development against genetic diseases linked to both NALCN- and SNARE protein-containing complexes.

## Introduction

The resting membrane potential of neurons and other excitable cells is set by the push and pull of two opposing constitutively active ion fluxes – K^+^ exiting the cell, and Na^+^ entering. Whereas the K^+^ efflux is primarily driven by a large family of two-pore potassium (K2P) channels^1^, there is a sole molecular cause for up to 70% of the tonic Na^+^ influx – the sodium leak channel NALCN^2^. The Na^+^ influx carried by NALCN plays a key role in a wide range of physiological behaviours from respiratory rhythm to locomotion, and NALCN channelopathies result in severe neurodevelopmental disorders^3,4^.

Early challenges in the heterologous characterisation of this unusual relative of voltage-gated sodium and calcium channels were resolved by identifying three mandatory auxiliary subunits: UNC79, UNC80, and FAM155A, without which the channel is non-functional^5^. Together, these four proteins form a structurally unique channel complex with a large intracellular surface potentially capable of hosting other protein-protein interactions (Figure 1A)^6–8^. A range of endogenous proteins have been suggested to functionally and physically interact with the NALCN core complex, including a number of different G protein coupled receptors^9–11^, Src-family kinases^9,12^, and the Slo2.1 ion channel^13^. In addition, calmodulin has been shown to copurify with the channel complex^6,8^, although the functional implications of this direct interaction remain unclear.

**Figure 1:**
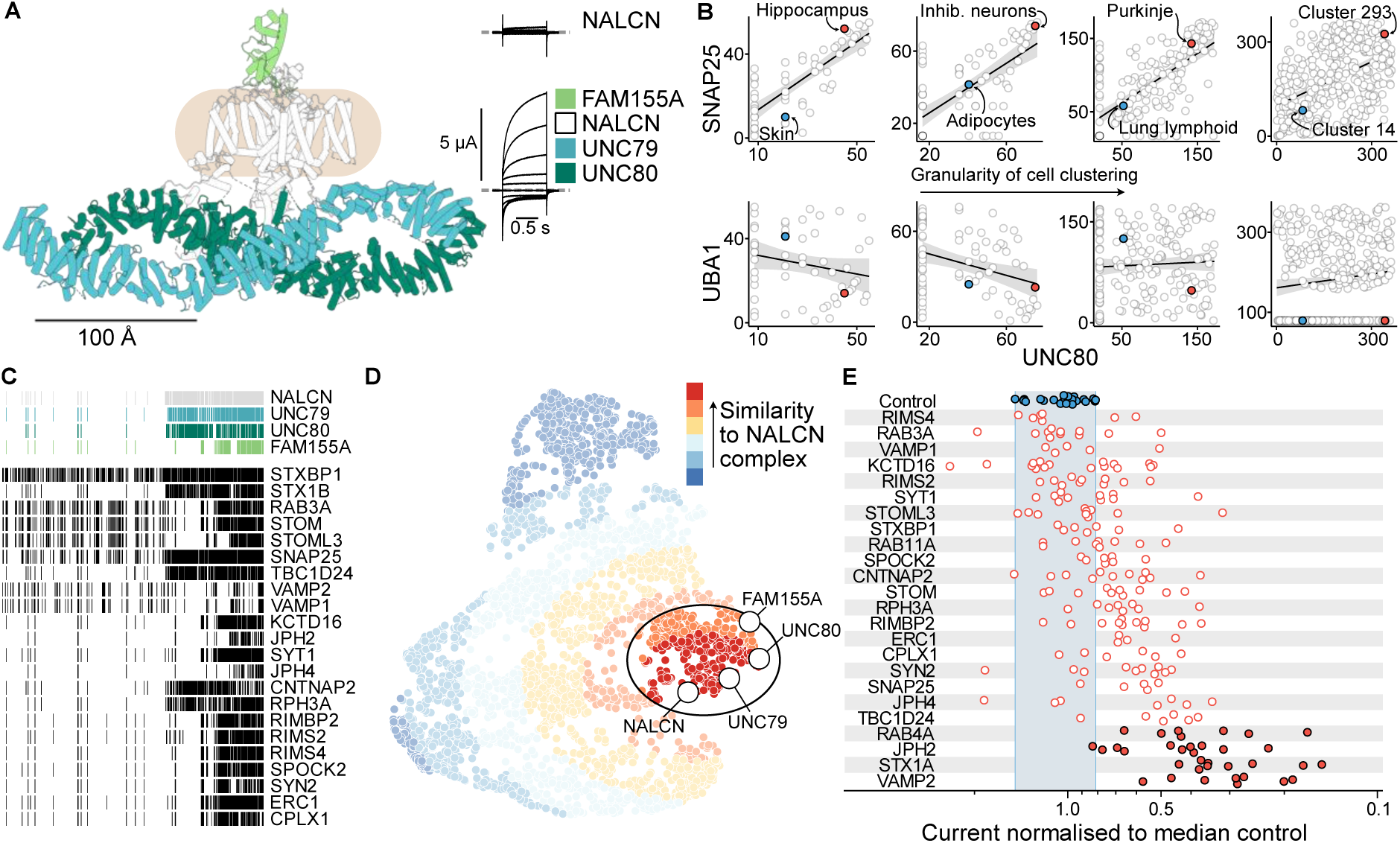
A computational screen identifies potential protein partners of the NALCN core complex. A) Cartoon representation of the NALCN core complex from PDB #7SX3. The location of the nanodisc is indicated by a light brown oval. Example current traces from Xenopus oocytes expressing NALCN alone (left) or in combination with the auxiliary subunits UNC79, UNC80 and FAM155A (right). Grey dashed lines indicate the zero current level. B) Correlations between UNC80 transcript expression ranks and either SNAP25 (top row) or UBA1 (bottom row) from HPA tissues, HPA cell types, the descartes atlas, and the Allen brain atlas (left to right). Two tissues/cell types are highlighted in each panel for comparison, with a neuronal type in red and a non-neuronal in blue. C) Patterns of ortholog presence and absence according to OrthoDB for the NALCN core complex and the genes chosen for functional screening, with each ‘barcode’ stripe representing the presence of an ortholog in a species. D) UMAP representation of the combined co-evolution and co-expression scores with each gene shown as a point. Proximity in UMAP space to the NALCN core complex is shown by the colour legend. The genes chosen for functional characterisation were chosen from the cluster highlighted inside the black oval. E) Normalised Ca^2+^-sensitive inward current magnitudes. The four candidates with the strongest effects on NALCN currents chosen for follow-up experiments are highlighted as red filled points.

The regulation of the NALCN core complex by protein partners is of pressing interest for several reasons. Firstly, the channel is notorious for its lack of specific pharmacological agents, both endogenous and otherwise^3,14,15^. It is therefore tempting to hypothesise that interfering with the modulation of NALCN activity mediated by protein interactions such as those described above may offer attractive opportunities for pharmacological intervention and treatment of NALCN related disorders. Secondly, the subcellular trafficking and localisation of the core complex remains unresolved. It is unclear where, when and how the channel is delivered to and active at the membrane of the neurons it has been identified in. A clearer picture of the protein interactome of the channel will offer direct insight towards its position and role within a neuron and may offer clues to how it shapes local excitability.

To identify other proteins that may regulate NALCN function, we developed an integrative computational approach utilizing i) co-expression of coding RNA transcripts, ii) co-evolution of gene orthologs, iii) shared gene-phenotype associations. This approach suggested a potential interaction with synaptic function-associated proteins. We found proteins of the neuronal SNARE complex inhibit NALCN complex function in a heterologous expression system, and structurally associate with the intracellular linker regions of NALCN. Furthermore, this interaction can be disrupted in primary neurons to increase the sodium leak, leading to rapid cell death. Together, these findings suggest a novel role for SNARE complex proteins in regulating the sodium leak current in neurons.

## Results

### A computational screen identifies putative protein partners of the NALCN core complex

We expected that proteins which interact with the NALCN core complex would be required to be expressed in the same cells. To identify proteins that share similar expression patterns to the four proteins that constitute the NALCN core complex, we calculated pairwise correlations between RNA transcripts coding for each protein-coding gene against each NALCN complex protein from bulk RNA sequencing of different human tissues^16^, or pseudobulk RNA sequencing from cell types in humans^16,17^ and mice^18^. We find that the presence of RNA encoding for NALCN complex proteins is highest in neuronal and other excitable cells and correlates well with each other (Figure S1A) and other proteins involved in neural function. For example, we see a positive correlation between UNC80 and SNAP25 transcript expression across bulk tissue and increasingly granular cell type clustering (median Spearman’s rank correlation coefficient 0.59), whereas there is little correlation between transcript expression of UNC80 and the cell-essential E1 ubiquitin-activating enzyme UBA1^19^ (Figure 1B, median Spearman’s rank correlation coefficient -0.03). Extending this analysis, we compared the differences in correlations between the NALCN core complex and gene sets i) linked to neuronal function^20,21^ (411 genes) and ii) essential for cell function^19^ (291 genes, Figure S1B, see Methods). We find more positive correlations between the NALCN core complex genes and neuronal genes than between the NALCN core complex genes and essential genes in 3 of the 4 datasets (median Spearman’s rank correlation coefficient of 0.24 and -0.22 respectively).

Our second approach is built on the assumption that proteins which function together or structurally interact experience similar evolutionary pressures, and so are likely to share a similar pattern of presence and absence across the tree of life^22,23^. We calculated fingerprint identity scores between the phylogenetic profiles of each human protein-coding gene and each of the four NALCN core complex genes based on orthology assignments of OMA^24^, OrthoDB^25^, and eggNOG^26^ for over 2000 unique eukaryotic species (Figure 1C). We saw nearly identical (88.3%, 99.5%, and 99.4% identity from each source respectively) phylogenetic profiles for UNC79 and UNC80 from all three sources of ortholog assignments, with the similarity of profiles for NALCN and FAM155A varying between sources (Figure S2A).

Finally, we restricted our selection to genes with genetic associations to clinically observed phenotypes similar to those reported for NALCN core complex genes. To do so, we used the Open Targets Platform^27^, which compiles data from a range of sources to produce a disease-association score for a given gene based on the weighted strength of evidence. We selected a longlist of 2554 genes with disease-association scores above a threshold of 0.01 for one or more of the general phenotypes ‘Intellectual disability’, ‘Abnormal pattern of respiration’, ‘Neurodevelopmental disorder’, or ’Arthrogryposis syndrome’, all of which are broader categories used to describe the symptoms which occur in the NALCN-related channelopathies CLIFAHHD (Congenital Contractures of Limbs and Face, Hypotonia, and Developmental Delay) and IHPRF1 (Infantile Hypotonia with Psychomotor Retardation and characteristic Facies 1)^3^.

We then projected the RNA co-expression and gene co-evolution scores for each longlisted gene into two dimensions with UMAP (Figure 1D) and calculated the Euclidean distance between each individual gene and the centroid of the four NALCN core complex genes. Our shortlist of the most proximal 200 genes were more likely to be involved in molecular functions like synaptic assembly and less likely to be involved in metabolic or biosynthetic processes than the longlist (Figure S3A). Of the genes which have been previously suggested to modulate NALCN function, four appear in the disease-associated longlist (muscarinic acetylcholine receptors, calcium-sensing receptor, tachykinin receptor 1, calmodulins) but we only found genes encoding muscarinic acetylcholine receptor subtypes (CHRM1/2) in the most proximal 200 gene shortlist.

### Four candidate proteins inhibit currents of the heterologously expressed NALCN core complex

Guided by the over-representation of synaptic function-related genes in the shortlist, we predominantly chose proteins from the list that were known to be involved in synaptic function (e.g. STX1B, SNAP25, CPLX1). In addition, we chose proteins from the list that have also been previously suggested to regulate other ion channels (e.g. KCTD16^28^, JPH2^29^, RAB3A^30^). We also included additional isoforms of some of the chosen proteins that have also been linked to ion channel function (RAB4A^30^, RAB11A^30^, STX1A^31^, CPLX2^32^). As a result, we continued with 25 proteins for functional screening.

To screen for functional effects on NALCN core complex function, we injected mRNA coding for each candidate protein one day after injecting a mix of mRNAs coding for NALCN, UNC79, UNC80 and FAM155A in *Xenopus laevis* oocytes. Five days after the initial injection, we recorded sodium leak currents by two-electrode voltage clamp (TEVC). We took advantage of the strong inhibition of inward NALCN-mediated currents by extracellular Ca^2+^ and Mg^2+^ to isolate NALCN complex-specific activity^5^. We took the initial Ca^2+^-sensitive inward current in response to a hyperpolarising voltage step as the NALCN complex-specific functional readout (Figure S3B). We found four candidate proteins in particular exhibited strong inhibitory effects on NALCN function: RAB4A, JPH2, STX1A, and VAMP2 (Figure 1E).

Many of the tested candidate proteins are known to interact with each other, and in some cases form macromolecular complexes. To discern whether these interactions were required for functional effects on NALCN, we injected combinations of mRNAs coding for candidate proteins which have been shown previously to interact (Figure S3D). Notably, we found that injecting a combination of STX1A and SNAP25, two components of the neuronal SNARE (Soluble N-ethylmaleimide-sensitive factor attachment protein receptors) complex (Figure 2A), leads to stronger inhibition of NALCN currents than STX1A alone, while SNAP25 alone had no discernible effect (Figure 2B-C). Although we saw inhibition of NALCN currents by high concentrations of VAMP2 (the third member of the tripartite core neuronal SNARE complex) alone, we did not see inhibition of NALCN currents at lower concentrations, or additional potentiation of inhibition in combination with STX1A or SNAP25 (Figure 2C).

**Figure 2:**
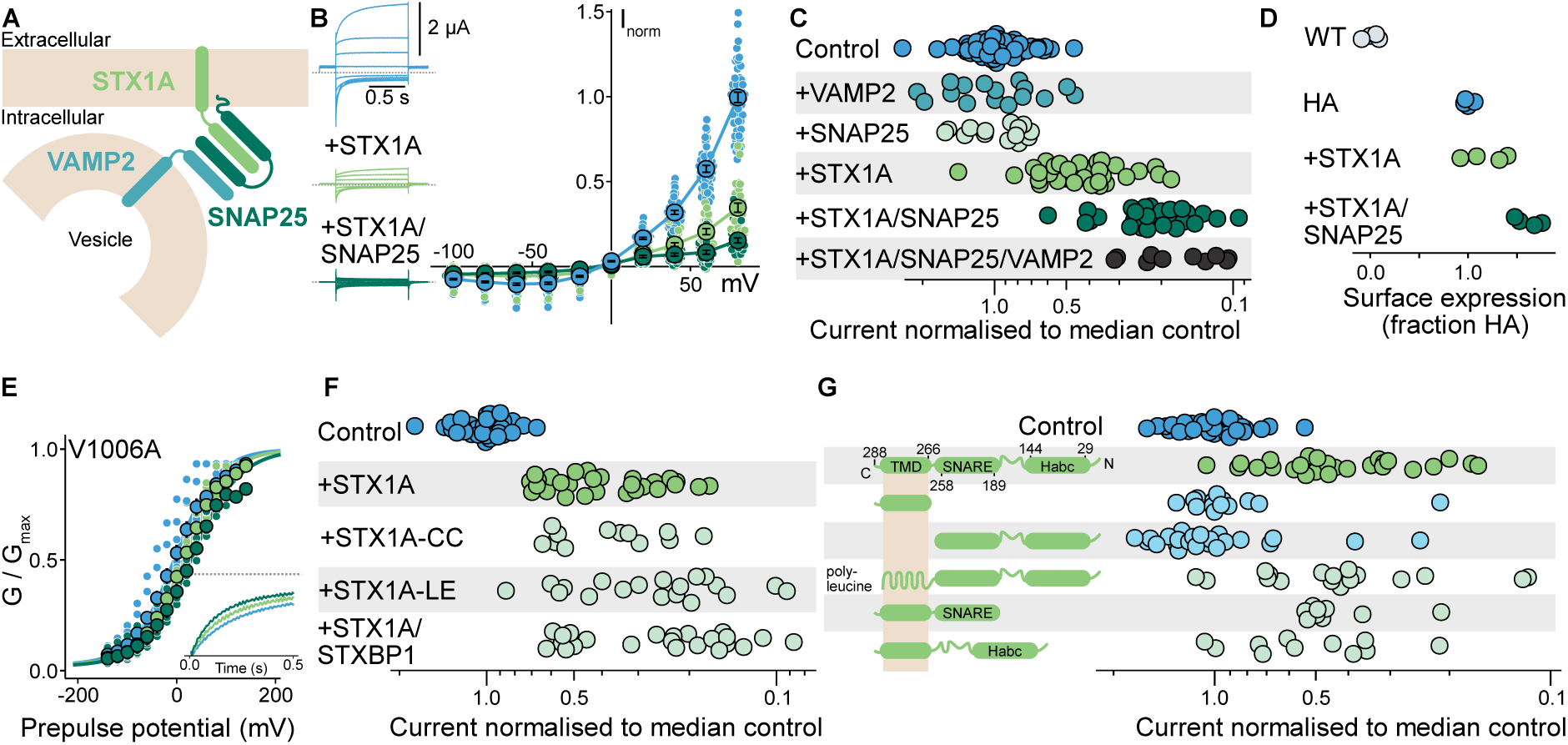
Neuronal SNARE complex proteins reduce NALCN activity in vitro. A) Cartoon overview of the SNARE protein complex assembly. B) Left: representative current traces from Xenopus oocytes expressing the NALCN core complex alone (blue), or additionally expressing STX1A or STX1A-SNAP25. Right: Normalised steady-state current-voltage relationships for the same conditions as on the left. C) Normalised inward current magnitudes at -100 mV from oocytes expressing different combinations of SNARE proteins with the NALCN core complex. D) Normalised surface expression of the NALCN core complex +/- STX1A-SNAP25. E) Conductance-voltage curves from Xenopus oocytes expressing the NALCN core complex alone (blue), or additionally expressing STX1A (light green) or STX1A-SNAP25 (dark green). Inset: representative current deactivation traces for NALCN-V1006A expressed either with UNC79-UNC80-FAM155A alone (blue), or additionally expressing STX1A (light green) or STX1A-SNAP25 (dark green). F) Normalised inward current magnitudes from cells expressing the NALCN core complex alone or with variants of STX1A. G) Left: cartoon overview of the topology of the different STX1A constructs tested (see SI for construct sequences). Right: normalised inward current magnitudes from cells expressing the NALCN core complex alone or in combination with the STX1A constructs shown in the left panel.

Due to the striking inhibitory effect of STX1A on NALCN currents and the intriguing potentiation of said inhibition by fellow SNARE complex component SNAP25, we decided to focus our efforts on characterising the functional, structural, and physiological ramifications of the STX1A-SNAP25 inhibition of sodium leak currents.

### Neuronal SNARE complex proteins reduce NALCN activity in vitro

Together, STX1A, SNAP25 and VAMP2 form the core of the neuronal SNARE complex, the protein assembly that drives the membrane fusion events required for neurotransmitter release at the synapse^33^. Each of the three proteins is a member of a wider family of SNAREs (over 60 members in mammals^34^) which drive the fusion of vesicles and membranes more generally across cells^33,35^.

The inhibition of NALCN currents observed from co-expression of SNARE complex proteins may result from two broad mechanisms: lower channel activity, and/or lower channel expression at the cell membrane. To determine whether STX1A and SNAP25 altered the trafficking of NALCN to the cell membrane, we introduced an HA-epitope into an extracellular loop of NALCN and measured surface expression in *Xenopus* oocytes (Figure 2D). The introduction of the HA-epitope does not alter NALCN core complex assembly or function (Figure S4). We find that neither STX1A alone nor STX1A-SNAP25 together reduce the surface expression of NALCN at the membrane (Figure 2D). Together, these results suggest the inhibition of NALCN currents by STX1A-SNAP25 is mediated by reducing channel activity, not channel surface expression.

NALCN channel activity can be modulated through several mechanisms, such as altering the open probability or changing the characteristics of its response to changes in voltage. We investigated the effects of co-expressing STX1A and SNAP25 on the voltage-sensitivity of activation and deactivation of NALCN currents. While a leak channel by nature, NALCN complexes do exhibit shallow conductance-voltage relationships with a voltage of half-activation (V_mid_) estimated here of around +50 mV. For wild-type channels, we were unable to achieve fully saturating prepulse voltages while maintaining the integrity of the cell membrane (Figure S5A). We therefore introduced two separate gain-of-function mutations (V1006A or R1181G) into NALCN^14^, which reduce the V_mid_ of the channel to more negative potentials (Figure 2E, S5A,B). Co-expression of STX1A and STX1A-SNAP25 with either of these point mutants shifts the V_mid_ slightly towards more positive potentials, with the combination of STX1A-SNAP25 having a larger effect. We did not see a noticeable modulation of the voltage-insensitive component of NALCN current, which is estimated here at 2-5% of the maximal channel conductance. Additionally, we observe that the rate of deactivation of inward current in response to a hyperpolarising voltage step was somewhat increased by STX1A and STX1A-SNAP25 for both mutant channels (Figure 2E inset & Figure S5B).

In *Xenopus* oocytes, STX1A alone or together with SNAP25 does not reduce the amount of NALCN present in the plasma membrane. There is some evidence for modulation of the voltage-dependence of channel activity such that more positive membrane potentials are required for channel opening, and channels deactivate slightly faster on returning to more negative membrane potentials. However, these biophysical effects are small and are unable to fully explain the dramatic reduction in macroscopic currents we observe. We hypothesize that in addition to these small effects, there is a dramatic reduction in the ability of the channel to open across all voltages which underpins the loss of macroscopic current which we observe.

### STX1A requires membrane localization to inhibit NALCN core complex activity

STX1A exists predominantly in one of two conformations: a self-inhibited ‘closed’ conformation in which the Habc domain folds back onto the SNARE domain, and an ‘open’ conformation that allows for SNARE complex assembly and membrane fusion^36–38^. To determine whether the conformational state of STX1A is important for its inhibition of NALCN complex currents, we expressed a construct with two residues in the ‘hinge’ region of the protein between the SNARE and Habc domains (STX1A-LE, L165A and E166A) mutated to alanine. This STX1A-LE construct is mostly found in the open state, with the SNARE and Habc domains dissociated^36^. We also co-expressed STX1A with STXBP1, a protein which interacts with STX1A to favour its closed state^37,39^. Biasing STX1A towards either its closed or open conformations has no discernible effect on its inhibitory effect on NALCN complex currents (Figure 2F). Similarly, substitution of two cysteines in the transmembrane domain of STX1A that were previously implicated as crucial for its interactions with the Ca_V_1 and Ca_V_2.2 channels (STX1A-CC, C271V and C272V)^40^ does not abolish the effect on NALCN as it does for Ca_V_s (Figure 2F).

Next, we attempted to narrow down the interacting domains of NALCN and STX1A by generating a series of STX1A truncations. STX1A constructs missing either the SNARE (residues 189-258) or Habc (residues 29-144) domains were still able to inhibit NALCN currents, but when the entire intracellular domain of the protein was removed to leave only the transmembrane segment, the inhibitory effect was lost (Figure 2G). However, when we expressed the intracellular domain in isolation, we saw no STX1A-like inhibition. Inhibition could be restored by re-localizing the intracellular domain to the membrane with a polyleucine helix. Together, our data highlight the necessity of STX1A’s localisation to the cell membrane for its inhibition of NALCN.

### STX1A-SNAP25 interact with the DII-DIII linker of NALCN

To explore whether the functional effects observed in oocytes were the result of a direct interaction, we co-expressed the NALCN core complex with STX1A and SNAP25) in Expi293 cells and purified the resulting complex in detergent. Both STX1A and SNAP25 purified with NALCN and co-eluted during the final size exclusion chromatography (SEC) step (Figure S6A), suggesting the formation of a stable interaction between the neuronal complex proteins and NALCN. The sub-stoichiometric appearance of the intracellular UNC79 and UNC80 subunits relative to NALCN, STX1A and SNAP25 suggests that UNC79 and UNC80 are not required for the interaction of STX1A and SNAP25 (Figure S6A). Indeed, co-expression of only NALCN and FAM155A with STX1A and SNAP25 was sufficient for the purification of a stable NALCN-FAM155A-STX1A-SNAP25 complex (Figure 3A).

**Figure 3:**
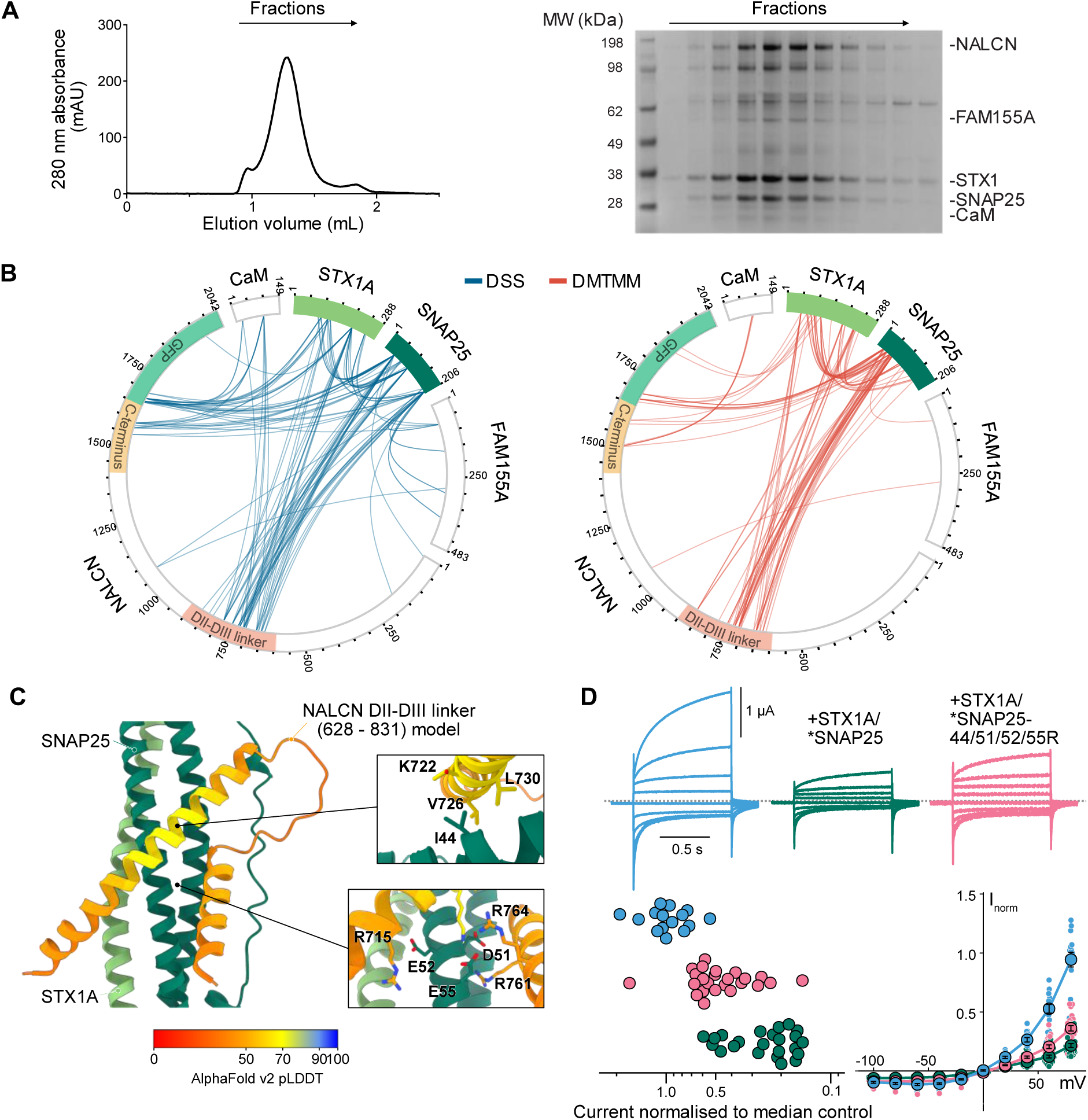
STX1A-SNAP25 interact with the DII-DIII linker of NALCN. A) Size exclusion chromatography (left) and corresponding SDS-PAGE (right) of the purified NALCN-FAM155A-STX1A-SNAP25 complex. B) Crosslinking mass spectrometry mapping of the protein-protein interactions between the complex members with DSS (disuccinimidyl suberate, left, blue lines) or DMTMM (4-(4,6-dimethoxy1,3,5-triazin-2-yl)-4-methylmorpholinium chloride, right, red lines). Selected regions of NALCN are highlighted in the outer circle C) Predicted interaction interface of the NALCN-DII-DIII linker and SNAP25/STX1A by AlphaFold v2 multimer. Highlighted interaction hotspots are shown with key residues on SNAP25 and the NALCN model labelled. D) Top: representative current traces from Xenopus oocytes expressing NALCN/UNC79/UNC80/FAM155A alone (blue), or additionally expressing Flag-tagged STX1A and SNAP25 (dark green) or STX1A and SNAP25 harbouring four point mutations (pink). Grey dashed lines indicate the zero current level. Lower left: normalised current magnitudes from cells expressing the same constructs as above. Lower right: Steady-state current-voltage relationships from the same experiments.

To further dissect this structural interaction, we performed crosslinking mass spectrometry on the purified NALCN-FAM155A-STX1A-SNAP25 complex^41^. For this purpose, we used two orthogonal crosslinking reagents in parallel experiments: DMS (disuccinimidyl suberate), which primarily forms crosslinks between lysine residues; and DMTMM (4-(4,6-dimethoxy1,3,5-triazin-2-yl)-4-methylmorpholinium chloride), which connects lysine residues with aspartic or glutamic acid residues. The two chemistries are expected to form covalent bonds between reactive sites in close proximity (up to around 30 Å relative to the protein backbone)^41,42^. The location of these crosslinks can be recovered after protease digestion and mass spectrometry of the resulting fragments, with crosslinked peptides revealing structural details of the original protein complex. As previously described, we found endogenous calmodulin copurified and formed crosslinks with the C-terminal domain of NALCN^6^. We found abundant crosslinks formed between NALCN, STX1A, and SNAP25 (Figure 3B, Table S1) that were mutually supportive in experiments using two distinct crosslinking chemistries. The crosslinks suggest that both the intracellular DII-DIII linker and C-terminal tail of NALCN are in close proximity to STX1A-SNAP25. We were particularly intrigued by the crosslinks with the DII-DIII linker for two reasons. Firstly, this linker has been previously shown to form an elaborate clamp onto UNC80^6^. Given that the interaction can occur in the absence of UNC80 in the biochemical characterisation (Figure S6A), this raises the possibility that STX1A-SNAP25 is able to displace UNC80 by binding to the DII-DIII linker of NALCN. Secondly, the DII-DIII linker of NALCN corresponds to the ‘synprint (synaptic protein interaction) site’ of the voltage-gated calcium channels which have been previously been shown to bind to and be regulated by STX1A-SNAP25^31,43,44^, although they share very little sequence similarity with NALCN in this region.

Despite the stability of the purified NALCN-FAM155A-STX1A-SNAP25 complex, we were unable to obtain an experimental co-structure of the complex. Instead, we used AlphaFold v2 multimer^45^ to generate a predicted multimolecular complex between STX1A, SNAP25 and the NALCN DII-III linker (Figure 3C, Figure S6B-D). The prediction suggests a potential interaction between residues D699 and S771 of the NALCN DII-DIII linker and a three helical bundle of STX1A-SNAP25, in agreement with the crosslinks. In this model, SNAP25 contributes the majority of residues contacting the NALCN DII-DIII linker with an extended electrostatic interface between SNAP25 D51, E52, E55 and NALCN R715, K722, R761, R764 and the hydrophobic residues SNAP25 I44 and NALCN V726 and L730 (Figure 3C). This interaction between NALCN and STX1A-SNAP25 would be incompatible with the DII-DIII linker – UNC80 interaction observed in cryo-EM structures (Figure S6E)^6–8^.

We next tested the interaction *in vitro* with a recombinantly purified STX1A-SNAP25 complex and a synthetic NALCN-DII-DIII linker (E698-N772) peptide by SEC. The NALCN-DII-DIII linker peptide co-eluted with the STX1A-SNAP25 complex and strongly left-shifted the SEC elution profile compared to the elution of the STX1A-SNAP25 only (Figure S7). Of note, the NALCN DII-DIII linker-STX1A-SNAP25 complex on the SEC column eluted earlier than expected for a stoichiometric 1:1:1 (NALCN DII-DIII linker:STX1A:SNAP25) complex and suggests the formation of higher-order assemblies.

Mutating the four residues of SNAP25 which form the predicted interface (SNAP25-44/51/52/55R) resulted in no co-elution and shift on the SEC elution profile after incubating with STX1A and the NALCN-DII-DIII linker (E698-N772) peptide (Figure S7). Replacement of the four residues on SNAP25 resulted in weaker potentiation of STX1A inhibition of NALCN currents in oocytes (Figure 3D), restoring NALCN function to levels resembling co-expression with STX1A alone.

Taken together, the biochemical and electrophysiological data suggest that the inhibition of sodium leak currents by STX1A-SNAP25 is primarily mediated through an interaction with the intracellular DII-DIII linker of NALCN. We hypothesize that this interaction results in a change in the complex that prevents the channel from opening, given the lack of effects on channel surface expression and small changes in other biophysical parameters. An attractive mechanistic explanation for this dramatic reduction in channel activity is the displacement of UNC80 from the DII-DIII linker of NALCN by competition with STX1A-SNAP25, thus reducing the number of channels able to open. Curiously, despite the strong functional inhibition we observe from expression of STX1A alone, and the converging insights from crosslinking data, AlphaFold predictions, and truncation experiments, future studies will be required to elucidate the precise molecular nature of this interaction.

### STX1A-SNAP25 do not inhibit voltage-gated sodium channels

We were surprised by the shared STX1A-SNAP25 binding domain between NALCN and voltage-gated calcium channels, as the DII-DIII linker is not well conserved between the proteins. We wondered whether the STX1A-SNAP25 inhibition could be a conserved feature across the whole four-domain voltage gated channel family in humans (Ca_V_s, Na_V_s and NALCN). We therefore tested for functional effects of coexpressing STX1A and STX1A-SNAP25 with three different members of the voltage-gated sodium channel family (Na_V_1.4, Na_V_1.5, and Na_V_1.7) in *Xenopus* oocytes, but did not observe alterations of voltage-sensitivity of activation or peak current amplitude (Figure S8). These data suggest that STX1A-SNAP25 are not generally regulating all four-domain voltage-gated channels but are selective in which channels they are able to functionally modulate.

### Loss of syntaxin increases sodium leak in murine isolated hippocampal neurons

The data presented so far have been derived from overexpression of each of the relevant proteins in heterologous expression systems. Next, we sought to establish whether the functional effects are of physiological relevance in the more complex cellular environment of a neuronal cell. As we observe that overexpressing STX1A-SNAP25 inhibits NALCN function, and that the SNAP25 effect is conditional on the presence of STX1A, we hypothesised that removing STX1A from neurons should lead to an increase in the sodium leak current. In neurons, removal of STX1A is functionally compensated by its paralog STX1B^37,46^. We determined that STX1B was also able to recapitulate the inhibition of NALCN currents in our heterologous system, both alone and in combination with SNAP25 (Figure S9). To completely remove both syntaxin isoforms, we turned to a previously established Stx1A/1B conditional double knockout (*Stx1B*^FL/FL^/*Stx1A* KO) mouse line, where *Stx1A* is globally knocked out and *Stx1B* is flanked by LoxP sites and can be removed by Cre recombinase expression^47^.

We isolated hippocampal neurons from newborn wild-type or *Stx1B*^FL/FL^/*Stx1A* KO mice and cultured them at high density on astrocyte feeder layers from wild-type mice. We recorded sodium leak currents from neurons between 14 and 22 days after plating (14-22 DIV) in response to lowering the concentration of extracellular divalent ions or replacing the extracellular sodium with NMDG (Figure 4A). We found that at -70 mV, wild-type neurons exhibited an increase in sodium leak of 0.25–0.41 pA/pF when the perfused concentration of Ca^2+^ and Mg^2+^ was lowered from 2 mM to 0.5 mM (Figure 4B), reflecting the release of inhibition of sodium leak currents by divalent ions^5^. This leak current was halved to 0.12–0.24 pA/pF by knocking down *Nalcn* expression by around 70% (Figure S11C) with a lentiviral short-hairpin RNA construct (shNalcn) at DIV1, suggesting that the sodium leak current under these recording conditions is predominantly conducted by Nalcn. These findings are in line with those reported by others, with the NALCN-mediated contribution to sodium leak estimated at between 60% to 80% in murine hippocampal^2^, SCN^48^, and midbrain dopaminergic neurons^10^.

**Figure 4:**
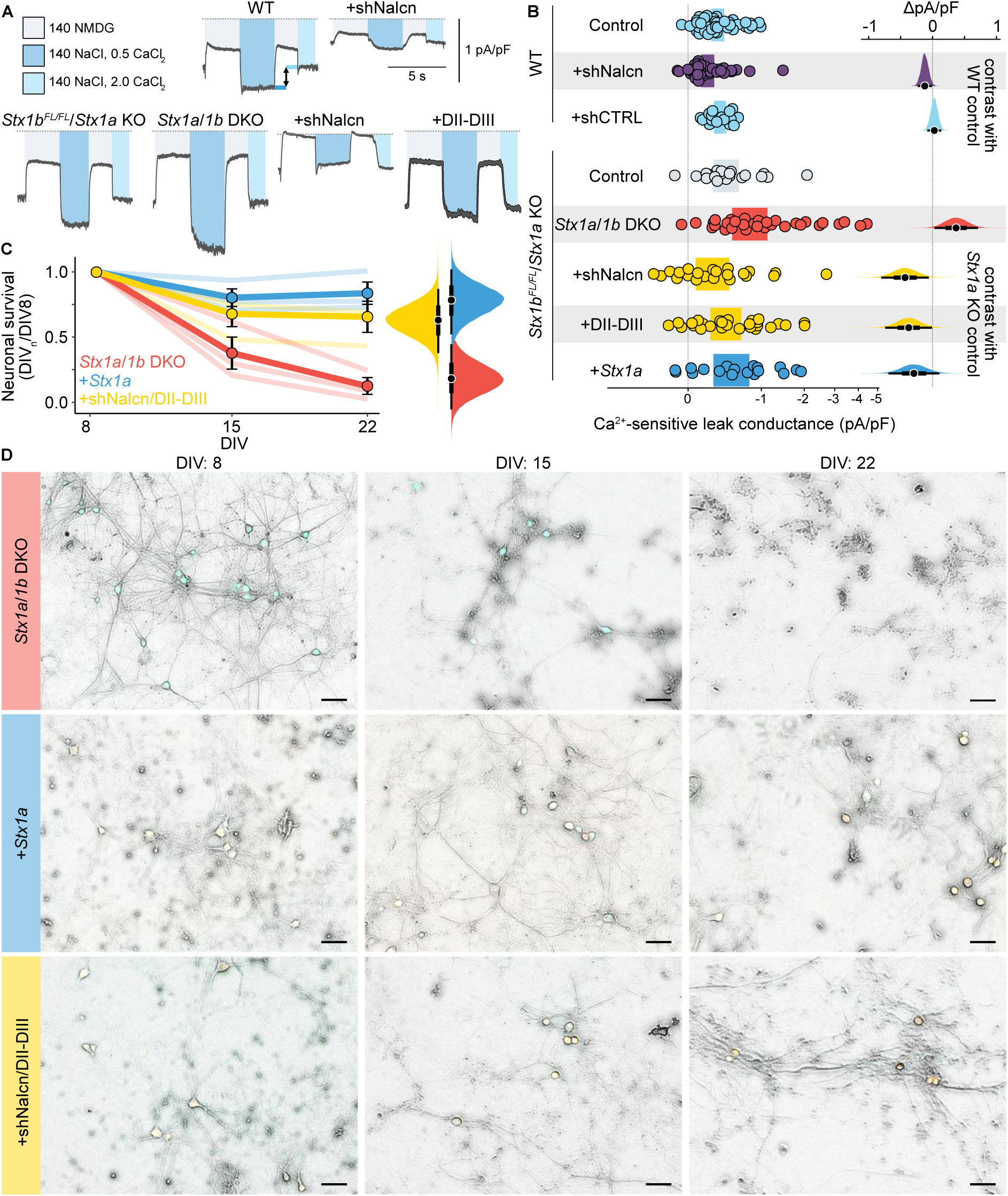
Loss of syntaxin increases sodium leak in isolated hippocampal neurons. A) Example current traces from cultured hippocampal neurons held at -70 mV and perfused with the indicated extracellular solutions (full solution compositions are in the Methods). The Ca^2+^- sensitive leak conductance is calculated as shown by the arrow in the example trace for WT neurons B) Left: summary Ca^2+^-sensitive holding current magnitudes for neurons isolated from WT (top) or Stx1B^FL/FL^/Stx1A KO (bottom) mice infected with lentiviral constructs at DIV 1 as indicated. Individual data points are from 2 or more different primary cultures and recorded from DIV 14-22. The 95% credible intervals of the group-level estimates are shown as coloured blocks. The grey dashed line indicates the zero current level. Data is shown on a log2(1+x) scale. Right: contrasts calculated from the distributions of the expected value of the posterior are shown in full, with the median values shown as points and the 66% and 95% quantiles shown as thick and thin black lines, respectively. C) Fraction of plated hippocampal neurons isolated from Stx1B^FL/FL^/Stx1A KO and infected with the indicated lentiviral constructs at DIV 1 surviving in culture over time. Points and error bars represent the mean and standard error of the mean. Individual cultures are shown by translucent lines. The distributions of the expected value of the posterior for the surviving fraction of neurons at DIV 22 are shown in full to the right, with the median values shown as points and the 66% and 95% quantiles shown as thick and thin black lines, respectively. D) Representative merged brightfield and fluorescent images of cultured neurons from the same culture from DIV 8 to DIV 22. Scale bars are 50 µm. Contrast adjustment and background correction were performed for clarity here; uncorrected images can be found in Figure S12.

Non-infected *Stx1B*^FL/FL^/*Stx1A* KO neurons, which lack Stx1A but still express Stx1B, exhibited sodium leak currents of similar magnitude to wild-type neurons, consistent with our observations in *Xenopus* oocytes that STX1B/SNAP25 are also capable of reducing NALCN currents (Figure S9). Strikingly, removing Stx1B expression by lentiviral expression of Cre recombinase resulting in *Stx1A*/*1B* DKO neurons led to an increase in sodium leak to 0.52 - 1.13 pA/pF. Knockdown of *Nalcn* expression with shNalcn or inhibition of Nalcn currents by inducing expression of an interfering protein (DII-DIII, corresponding to residues 617-835 of human NALCN) reduced sodium leak currents to within the range of wild-type neurons, although with considerable variability (Figure 4B). This interfering peptide disrupts the formation of the NALCN complex by competing for the DII-DIII linker interaction site on UNC80, dramatically reducing sodium leak currents^6^. Together, these data demonstrate that sodium leak currents mediated by Nalcn in hippocampal neurons are inhibited by Stx1a/1b, and removal of these proteins enhances the NALCN-mediated sodium leak.

### Reducing sodium leak currents enhances survival in neurons lacking both syntaxin isoforms

In addition to the role of Stx1A/1B in supporting neurotransmitter release, *in vitro* genetic removal of both isoforms leads to neuronal cell death (Figure 4C,D) as shown previously^47,49,50^. Curiously, the ability of Stx1A/1B to support neurotransmitter release is independent from its ability to keep neurons alive, as reintroducing fusion-incompetent Stx1a isoforms into neurons lacking endogenous Stx1A/1B prevents cell death^47^, suggesting an as-yet unexplained role for Stx1A/1B in neuronal survival. We were curious to test if the increase in sodium leak currents resulting from removal of Stx1A/1B could represent the possible mechanism underlying the resulting cell death. We found that while reducing Nalcn expression (shNalcn) or currents (DII-DIII) individually did not result in improved survival (Figure S11B, Figure S12), combining these two interventions to reduce sodium leak currents enhanced neuronal survival such that around half of *Stx1A*/*1B* DKO neurons present at DIV8 were still alive at DIV22 (Figure 4C). Although the morphology of the surviving neurons was still altered when compared to *Stx1A*/*1B* DKO neurons with *Stx1A* reintroduced by lentiviral expression (Figure 4D), these data clearly suggest a cytotoxic effect of NALCN-mediated sodium leak currents when inhibition by neuronal SNARE complex proteins is relieved.

## Discussion

In this study, we demonstrate that the SNARE complex proteins STX1A and SNAP25 inhibit NALCN currents in heterologous expression systems and mammalian neurons. Removal of endogenous Stx1A/B from isolated murine hippocampal neurons results in an increase in sodium leak currents and rapid cell death, and knockdown or inhibition of Nalcn currents mostly reverses both phenotypes *in vitro*. It is commonly accepted that the neuronal SNARE complex feeds back on the Ca^2+^-channels responsible for the Ca^2+^ influx required for SNARE activation. Our data suggest that SNARE complex proteins STX1A and SNAP25 are also able to regulate the background Na^+^ influx through NALCN, and thus modulate cellular excitability directly.

We discovered this interaction through an integrative computational approach combining clues from phylogenetic profiling, RNA sequencing datasets, and links to disease phenotypes. Similar approaches have been previously successful in identifying protein constituents of the mitochondrial calcium uniporter^51^ and pairing endogenous peptide ligands to their GPCR partners^23^. We screened only a small selection of proteins from our shortlist (25 of 200) here to determine how useful the approach could be for identifying new partners of macromolecular ion channel complexes. This validation supports the effectiveness of our integrative approach and highlights the potential to uncover novel protein interactions. We believe that future efforts could further be elaborated by predictive biophysical interaction techniques and more sophisticated machine learning approaches that could be promising directions; both for other ion channel complexes and for further picking apart the possible interactome of the NALCN channel complex.

A common concern of protein-protein interaction screening in heterologous systems, and a potential limitation of this study, is that overexpression – and therefore artificially high concentrations or molar ratios - of the component proteins can lead to structural interactions and functional effects that are unlikely to occur at more physiological conditions. However, as STX1A and SNAP25 are two of the most abundant proteins in mammalian neurons - especially in synaptic regions^52^ (estimated at over 20,000 copies of STX1A per synaptic bouton compared to e.g. ∼15 copies of Ca_V_2.1) – we expect the expression ratio of complex proteins to NALCN protein to be far higher in neurons than we achieve heterologously. In addition, the functional effects we observe in primary neurons are as a result of artificially depleting endogenous protein levels, and so are not subject to the same limitations.

SNARE complex proteins have been suggested to interact with a plethora of different ion channels: voltage-gated calcium channels^31,40,53–55^, voltage-gated potassium channels^56–62^, CFTR^63,64^, K_ATP_ channels^65,66^, ENaCs^67,68^; and modulate them by a variety of mechanisms, including alterations in protein trafficking, membrane expression, and biophysical parameters from voltage-sensitivity to single channel conductance levels. STX1A and SNAP25 have been shown to assemble in a diversity of hetero- and homo-oligomeric forms under different conditions^33^ and we cannot exclude that multimers of STX1A and/or SNAP25 play also a role in NALCN’s regulation. A key functional interaction appears to be mediated by the DII-DIII linker of NALCN to SNAP25A and STX1A (Figure 3B-D). Through its interaction with UNC80, the NALCN DII-DIII linker has previously been proposed as a nexus for NALCN channelsome assembly, gating and modulation^6^. A direct interaction of STX1A-SNAP25 with the DII-DIII linker and thereby competition with UNC80 may provide an intriguing mechanism for the inhibition of NALCN leak currents.

While the STX1A-SNAP25 complex is best known for its role at the presynaptic membrane where it coordinates neurotransmitter release, both proteins are ubiquitous across the neural membrane, including postsynaptic and extrasynaptic regions^69–71^. There is currently limited data on the subcellular localisation of NALCN core complexes in mammalian neurons^3^. It therefore remains unclear where in the neuron the functional interaction between the SNARE complex and the NALCN core complex may occur. The interaction between neuronal Ca_V_ channels and SNARE complex proteins has been suggested to play a key role in organising the intimate coordination of the neurotransmitter release machinery, ensuring that synaptic vesicles are held in close proximity to the calcium influx which then drives their fusion with the plasma membrane^38^. To our knowledge, there is little support to date for a synaptic localisation or role for sodium leak currents, and it is unclear how the NALCN core complex could be incorporated into this tightly organised subcellular environment. A definitive answer to this question would require the development of genetic or pharmacological tools to label and visualise endogenous NALCN complexes and their subcellular localisation.

An alternative proposal could be that STX1A-SNAP25 dimers that form outside the synaptic active zone (and comprise the bulk of SNARE proteins in mammalian neurons^72^) could regulate the NALCN complex in parallel to the neurotransmitter release cascade, thus diminishing sodium leak currents. The shared expression patterns of STX1A-SNAP25 and the NALCN core complex suggest this mechanism could occur widely across excitable cells. The most potent endogenous regulator of sodium leak currents described to date is extracellular calcium^5,73^, which can be reduced from its resting physiological level of around 1 mM (at which >80% of NALCN inward currents are inhibited^5^) by bursts of synaptic activity^74^ or during seizures^75^, relieving NALCN inhibition and contributing to increased cellular excitability. During these periods of greatest neuronal activity, STX1A-SNAP25 inhibition of the sodium leak current may be important for preventing excessive positive feedback and excitotoxicity, which may explain why genetic variation in both NALCN complex subunits and SNARE complex proteins can lead to seizures (discussed in more detail below).

One of the most surprising outcomes of this work is the contribution of sodium leak currents to neuronal cell death in the absence of Stx1A/B (Figure 4C-D). It has been shown by multiple groups that depletion of endogenous Stx1A/B in neurons leads to embryonic lethality in mice and widespread neuronal death^47,49,76,77^. Importantly, this essentiality for survival is independent of neurotransmission, as neuronal survival can be rescued by reintroducing an exocytosis-incompetent Stx1A mutant which eliminates synaptic transmission^47^. The molecular mechanism underpinning this role in neuronal survival is not understood, and even the cell death pathway triggered upon Stx1A/B depletion is atypical and does not resemble other established cell death processes in neurons^49^. Based on the results in this study, we propose that a missing link in the connection between Stx1A/B and neuronal survival is the sodium leak current conducted by the NALCN complex. Depletion of inhibitory Stx1A/B-SNAP25 proteins increases sodium leak currents and drives neuronal cell death. It remains unclear whether this cytotoxicity is directly caused by sodium influx, a more depolarised membrane potential, or other as-yet unexplored downstream signalling pathways.

Across the over 100 cases of NALCN/UNC79/UNC80 related genetic disorders (NALCN channelopathies) reported in the literature, there are a broad spectrum of clinical manifestations and severities^3^. Many of the common symptoms, including neurodevelopmental delay, intellectual disability, and seizures, are shared by patients with pathogenic mutations in neuronal SNARE genes (SNAREopathies)^78^. For both classes of disease, despite sharing a common pathogenic molecular origin, there is a strikingly diverse range of symptoms and clinical outcomes – even among different patients with mutations in the same gene. The underlying mechanisms behind this diversity are not fully understood, and could range from the level of functional redundancy for a given gene (e.g. FAM155B may compensate for loss of FAM155A function) to subtle differences in the expression and function of different complex components in different tissues and cell types^3,78^. The interaction between the two complexes we have uncovered here suggest another contributor to the symptomatic diversity, as SNAREopathies may result not just in altered synaptic transmission, but also altered cellular excitability and even neuronal survival through their modulatory role on the sodium leak current. A greater understanding of the interplay between these disease mechanisms may be beneficial for the development of future therapeutics for both NALCN channelopathies and SNAREopathies.

## Materials and Methods

### Computational screening

We used four RNA sequencing datasets to calculate gene expression correlations with the NALCN core complex: the Human Protein Atlas consensus tissue (accessed at: https://www.proteinatlas.org/download/rna_tissue_consensus.tsv.zip) and single cell type data (accessed at: https://www.proteinatlas.org/download/rna_single_cell_type.tsv.zip) datasets^16^, the Allen Brain Atlas mouse cortex and hippocampus 10x dataset (accessed at: https://portal.brain-map.org/atlases-and-data/rnaseq/mouse-whole-cortex-and-hippocampus-10x)^18^, and the descartes human fetal brain atlas (accessed at: https://descartes.brotmanbaty.org/bbi/human-gene-expression-during-development/)^17^. Gene-to-gene pairwise Spearmans rank correlations were calculated within each dataset between each of the NALCN core complex proteins (NALCN, UNC79, UNC80, FAM155A) and each human gene (or mouse ortholog). Mouse orthologs from the Allen Brain Atlas were mapped to their human orthologs using the Alliance of Genome Resources^79^.

We used three orthology databases - OMA^24^, OrthoDB^25^, and eggNOG^26^ - to calculate binary phylogenetic fingerprints for each human protein-coding gene across eukaryotic species, with 1 (positive bit) indicating the presence of an ortholog and 0 (negative bit) indicating the absence. We then calculated the fingerprint similarity score between each of the NALCN core complex protein-coding genes and each other human protein-coding gene as the number of identical bits divided by the total bits. Genes with low fingerprint similarity will have scores close to 0, while genes with high fingerprint similarity will have scores close to 1.

We compared the similarities to NALCN core complex components of both phylogenetic fingerprints and RNA expression profiles for a list of 411 genes annotated in the Gene Ontology database as being linked to neuronal function (evidence criteria limited to inferred from experiment, inferred from direct assay, inferred from mutant phenotype, author statement supported by traceable reference) and a list of 291 genes established as essential for cell survival^19^. This essential gene list has been previously used as a background control to identify protein-protein interactions through phylogenetic relationships^23^.

We used the OpenTargets platform^27^ to narrow our search space down to human genes associated with clinical symptoms similar to those that occur due to genetic variations in the NALCN core complex. We set a threshold for a combined evidence score of 0.01 for at least one of the general phenotypes ‘Intellectual disability’, ‘Abnormal pattern of respiration’, ‘Neurodevelopmental disorder’, or ’Arthrogryposis syndrome’.

From the 2554 genes which met this threshold, we created a gene by score matrix where each row is a human protein-coding gene associated with similar clinical phenotypes to NALCN channelopathies, and each column is a score between 0 and 1 representing either a Spearmans rank correlation with a NALCN core complex component’s RNA expression profile or a fingerprint similarity score with a NALCN core complex component’s phylogenetic fingerprint. The matrix was centred and normalised by subtracting the mean and dividing by the standard deviation for each column, and then projected into two dimensions using UMAP. As the exact units of the two dimensions are not directly interpretable, we have shown the plot without a scale. The 200 genes with the shortest Euclidean distance to the centroid of the NALCN core complex genes were subjected to statistical over-representation analysis of their Gene Ontology molecular function annotations using the PANTHER online tool (Figure S3A)^80,81^. 22 genes were picked from this shortlist of 200 for functional screening, with 3 additional isoforms of the 22 included for a total of 25 genes.

### Molecular biology for heterologous expression

Human NALCN, UNC79, UNC80 and FAM155A complementary DNAs (cDNAs), codon optimized for *Homo sapiens*, cloned into the pCDNA3.1(+) vector, were used as previously described^6^. All deletions, substitutions and insertions were generated using site-directed mutagenesis with custom-designed primers (Eurofins Genomics/Sigma-Aldrich) and the Q5 Hot Start High-Fidelity DNA Polymerase (New England Biolabs). cDNAs for each of the candidate interacting proteins were synthesised and cloned into the pCDNA3.1(+) vector commercially (Twist Bioscience). Sequences of cDNAs were verified by Sanger DNA sequencing and/or whole-plasmid sequencing (Eurofins). For expression in *Xenopus laevis* oocytes, cDNAs were linearized using XbaI and then transcribed to capped mRNAs with the T7 mMessage mMachine Kit (Ambion). To directly compare the biochemical and functional experiments with the quadruple mutant SNAP25-44/51/52/55R, we tested the functional effects of introducing FLAG-tags into STX1A and SNAP25 (Figure S10). The functional effects were not abolished by the introduction of the tags, albeit the magnitude of inhibition was slightly reduced when compared to wild-type protein. DNA and AA sequences of the constructs generated for TEVC are in Table S2.

### Two-electrode voltage-clamp electrophysiology

*Xenopus laevis* oocytes were prepared as previously described^6^. Healthy-looking stage V–VI oocytes were isolated and injected with 32–41 nl of mRNA using a Nanoliter 2010 or Nanoliter 2020 injector (World Precision Instruments). Unless otherwise stated, all mRNAs were injected at a final concentration of 250 ng/µL, so that experiments were performed with 1:1 by weight ratios of components. For the initial functional screen, putative interacting proteins were first injected at 1000ng/µL. We found that after reducing the injected amount of RNA of the four initial candidates, only STX1A still markedly inhibited NALCN currents at a 1:1 ratio by weight (i.e. around 5:1 molar weight ratio; Figure S3C). While the quantity of injected RNA does not correlate exactly with the quantity of expressed protein, we continued with the lower amounts of RNA to try and avoid artifacts from overloading the ribosomal machinery.

Injected cells were incubated in ND96 storage solution (96 mM NaCl, 2 mM KCl, 1 mM MgCl_2_, 1.8 mM CaCl_2_, 5 mM HEPES, 2.5 mM pyruvate and 0.5 mM theophylline; pH 7.4 with NaOH) supplemented with 50 µg/ml gentamycin and tetracycline at 18 °C at 140 rpm. Four to five days after RNA injection, two-electrode voltage-clamp (TEVC) measurements were performed on oocytes continuously perfused in one of two ND96-based recording solutions: ND96 (96 mM NaCl, 2 mM KCl, 1 mM MgCl_2_, 1.8 mM CaCl_2_ and 5 mM HEPES (pH 7.4) with NaOH) or ND96 w/o Ca^2+^ (96 mM NaCl, 2 mM KCl, 1.8 mM BaCl_2_ and 5 mM HEPES (pH 7.4) with NaOH) at room temperature using a Warner OC-725C Oocyte Clamp amplifier (Warner Instruments). Data were acquired using the pCLAMP 10 software (Molecular Devices) and a Digidata 1550 digitizer (Molecular Devices), sampled at 10 kHz. Electrical powerline interference was filtered with a Hum Bug 50/60 Hz Noise Eliminator (Quest Scientific). Traces were further filtered for display offline with a 1kHz lowpass FIR filter with order 30. Recording microelectrodes with resistances around 0.2–1.0 MΩ were pulled from borosilicate glass capillaries (Harvard Apparatus) using a P-1000 Flaming/Brown Micropipette Puller System (Sutter Instrument) and were filled with 3 M KCl.

All TEVC experiments presented were performed on a minimum of two separate batches of oocytes from different *Xenopus* frogs. Unless otherwise stated, recordings were performed in ND96 w/o Ca^2+^ holding at 0 mV and stepping between -100 mV and + 80 mV in 20 mV increments for 1 second. Currents for IV plots were calculated at steady-state, at the end of the 1 second voltage step. To compare current magnitudes, the peak initial inward current from stepping to -100 mV was taken. To control for batch-to-batch variability, we normalised both peak current magnitudes and IV plots to the median control (oocytes expressing NALCN-UNC79-UNC80-FAM155A alone) current recorded from the same oocyte batch. For current magnitude comparison plots, data from each oocyte is shown as a separate point. For IV plots, individual oocytes are represented by points with white outlines, with black outlines representing the mean and error bars the standard error of the mean.

For conductance-voltage recordings of NALCN, tail currents were first recorded in ND96 w/o Ca^2+^ by holding at 0 mV, stepping to a prepulse potential (-140 mV to + 140 mV in 20 mV increments) for 2.5 seconds, then holding at -80 mV to record a tail current. The protocol was repeated while perfusing NMDG96 - a solution with sodium replaced by the NALCN-impermeable ion NMDG^+^ (96 mM NMDG^+^, 2 mM KCl, 1.8 mM BaCl_2_ and 5 mM HEPES (pH 7.4) with HCl). Sodium-specific currents were calculated as the difference in inward tail currents between ND96 and NMDG96 (Figure S4A). The peak inward tail currents at each prepulse voltage for each oocyte were normalised to the maximum observed current. Conductance-voltage relationships for each construct were fit to a Boltzmann equation (Figure S4B, upper). Deactivation kinetics were fit from the same recordings by calculating the sodium-specific prepulse currents at -100 mV, normalising each trace to the initial inward current, and fitting a single exponential (Figure S4B, lower).

For Na_V_1.4, Na_V_1.5, and Na_V_1.7 recordings, mRNA was injected at 20 ng/µL two days before recording, with STX1A and SNAP25 injected on the same day at 250 ng/µL. Na_V_1.4 is the rat sequence, while Na_V_1.5 and Na_V_1.7 are the human sequences. Currents were recorded in ND96 with P/4 capacitance correction. Conductance was calculated from currents by estimating the reversal potential of Na^+^ ions from a linear fit to the current between 20 mV and 40 mV, then applying the following formula: G = I / (V_command_ – V_reversal_) and normalising to the maximum conductance observed.

### Surface expression assay in Xenopus oocytes

Surface expression was measured by an adaptation of a previous protocol^82^. NALCN-HA was generated by inserting an HA epitope plus short glycine linkers (GGYPYDVPDYAGG) between residues E232 and L233 in the extracellular loop immediately after S5 of domain I in NALCN. This modified channel was functionally indistinguishable from wild-type NALCN in *Xenopus* oocytes (Figure S4). Oocytes were injected and maintained as for TEVC experiments. Groups of 9-15 oocytes were pooled and surface labelled after 5 days by first blocking for 1 hour in ND96 with 1% BSA (Sigma-Aldrich), incubating with 1 µg /ml mouse monoclonal anti-HA antibody (2-2.2.14, Invitrogen) in ND96 with 1% BSA for 1 hour, washing three times with ND96 with 1% BSA, and incubating with 2 µg/mL HRP-conjugated goat anti-mouse IgG (G- 21040, Invitrogen) in ND96 with 1% BSA for 1 hour. All steps were performed at 4 °C. Oocytes were then washed three times with ND96 before luminescence was measured using a LUMIstar Omega microplate reader (BMG Labtech) with a 10 second integration time. Gain settings varied depending on the oocyte batch. Data are presented normalised to the raw luminescence values observed for NALCN-HA/UNC79/UNC80/FAM155A for the same batch of oocytes. Each point represents a single well of 9-15 oocytes from 2 or more batches of oocytes.

### Expression and purification of NALCN-STX1A-SNAP25 complexes

Optimized coding DNA for human NALCN, FAM155A, UNC80, UNC79, SNAP25, STX1A were each cloned into a pRK vector behind a CMV promoter. A GFP-Flag-Twin-Strep tag II was added to the C terminus of NALCN and a Flag tag was added to the C terminus of FAM155A, UNC80, UNC79, SNAP25 and STX1A. Untagged SNAP25 and STX1A were co-expressed and co-purified for the complexes used for crosslinking mass spectrometry. Expi293F cells (Thermo Fisher) in suspension were cultured in Expi293 Expression medium under 5% CO_2_ at 37 °C and transfected using the ExpiFectamine 293 transfection kit, as per standard manufacturer protocols, with all DNAs at used at equimolar ratio. Transfected cells were cultured for 48 h before collection.

For all purifications involving NALCN protein, cell pellet was resuspended in 1:5 (weight:volume) volume of lysis buffer (25 mM HEPES pH 7.5, 200 mM NaCl, 1 μg ml^−1^ benzonase, 1 mM PMSF and Roche protease inhibitor tablets). Cells were lysed by dounce homogenization and the NALCN complex was subsequently solubilized by addition of 2% (w/v) GDN supplemented with 0.1% (w/v) cholesteryl hemisuccinate and 0.2 mg ml^−1^ porcine brain polar lipid extract (Avanti) for 2 h at 4 °C under gentle agitation. Insoluble debris was pelleted by ultracentrifugation at 125,000*g*_max_ for 1 h, and the supernatant containing the solubilized protein was collected for affinity purification by batch-binding to 5 ml of M2-agarose FLAG resin (Sigma) for 1 h at 4 °C. Unbound proteins were washed with 6 column volumes (CV) of purification buffer A (6 CV 25 mM HEPES pH 7.5, 200 mM NaCl and 0.04% (w/v) GDN) and 10 CV buffer supplemented with 5 mM ATP and 10 mM MgCl_2_. NALCN was eluted with 5 CV of purification buffer supplemented with 300 μg ml^−1^ FLAG peptide (Sigma). The eluent was collected and applied to 3 ml Strep-Tactin XT high-affinity resin (IBA) and bound in batch for 2 h. Unbound proteins were washed with 10 CV of purification buffer A and eluted with 5 CV of purification buffer supplemented with 50 mM biotin. The NALCN complexes were then concentrated with an Amicon Ultra centrifugal filter device (100 kDa MWCO) concentrator to 5–10 mg ml^−1^ and applied to a Superose 6 3.2/300 column that had been pre-equilibrated in purification buffer B (25 mM HEPES pH 7.5, 150 mM NaCl and 0.01% (w/v) GDN). Peak fractions of the complexes were pooled and concentrated with an Amicon Ultra centrifugal filter device (100 kDa MWCO) and flash-frozen for storage.

For STX1A-SNAP25 wild-type and mutant complexes, cell pellet was resuspended in 1:5 (weight:volume) volume of lysis buffer (25 mM HEPES pH 7.5, 150 mM NaCl, 1 μg ml^−1^ benzonase, 1 mM PMSF and Roche protease inhibitor tablets). Cells were lysed by dounce homogenization and the NALCN complex was subsequently solubilized by addition of 2% (w/v) GDN supplemented with 0.1% (w/v) cholesteryl hemisuccinate for 2 h at 4 °C under gentle agitation. Unbound proteins were washed with 6 column volumes (CV) of purification buffer B (6 CV 25 mM HEPES pH 7.5, 150 mM NaCl and 0.04% (w/v) GDN). The protein was eluted with 5 CV of purification buffer supplemented with 300 μg ml^−1^ FLAG peptide (Sigma). The complexes were then concentrated with an Amicon Ultra centrifugal filter device (50 kDa MWCO) concentrator and applied to a Superose 6 10/300 GL column that had been pre-equilibrated in purification buffer B (25 mM HEPES pH 7.5, 150 mM NaCl and 0.01% (w/v) GDN). Peak fractions of the complexes were pooled and concentrated with an Amicon Ultra centrifugal filter device (50 kDa MWCO) and flash-frozen for storage.

### Chemical crosslinking and sample preparation for mass spectrometry

Samples were purified as above and provided at a protein concentration of 1.8 mg/ml in crosslinking buffer (25 mM HEPES pH 7.5, 200 mM NaCl, 0.01% GDN and 5% glycerol). For further processing, the concentration was adjusted to 1.0 mg/ml with the same buffer. Crosslinking was performed with a total of 50 μg protein per experiment, following previously described procedures^83,84^.

For crosslinking of primary amines, a 1:1 mixture of disuccinimidyl suberate (DSS)-d0 and d12 (Creative Molecules; from a 25 mM stock in dimethylformamide) was added to the sample to a final concentration of 1 mM and the sample was incubated for 30 min at 25 °C with mild shaking (750 rpm on the thermomixer). The reaction was stopped with 1 M ammonium bicarbonate (ABC) to a final concentration of 50 mM and was incubated for 20 min at 37 °C with mild shaking (750 rpm). The sample was dried by evaporation.

For crosslinking of carboxyl groups with primary amines, the sample was incubated with a 1:1 mixture of 22 mM pimelic dihydrazide (PDH)-d0 and d10 (ABCR;Sigma-Aldrich) together with 11 mM 4-(4,6-dimethoxy-1,3,5-triazin-2-yl)-4-methyl-morpholinium (DMTMM) chloride for 30 min at 25 °C. The reaction was stopped by passing the samples through a Zeba Spin Desalting column (7k MWCO, ThermoFisher Scientific). The filtrate was dried by evaporation in a vacuum centrifuge.

Dried samples were reconstituted in 8 M urea to a final concentration of 1.0 mg/ml and disulfide bonds were reduced by adding tris(2-carboxyethyl) phosphine to a final concentration of 2.5 mM. The samples were incubated for 30 min at 37 °C and cooled to room temperature prior to carbamidomethylation with iodoacetamide (IAA) added to a final concentration of 5 mM. Samples were incubated for 30 min in the dark and diluted with 150 mM ABC to a final concentration of ∼5.5 M urea; endopeptidase Lys-C (Wako) was added at an enzyme-to-substrate ratio of 1:100 and samples were further incubated for 2 h at 37 °C with mild shaking (750 rpm). The urea concentration was further diluted to 1 M using 50 mM ABC and trypsin (Promega) was added at an enzyme-to-substrate ratio of 1:50. Samples were incubated at 37 °C with mild shaking (750 rpm) overnight. On the next day, the samples were acidified by adding 100% formic acid to a final concentration of 2% and desalted by solid-phase extraction (Sep-Pak tC18 cartridges, Waters). The desalted samples were then fractionated by peptide-level size-exclusion chromatography (SEC) using a Superdex 30 Increase column (300 x 3.2 mm; GE Healthcare) (mobile phase: water/acetonitrile/trifluoroacetic acid (70:30/0.1, v/v/v), flow rate of 50 μl/min). Four 100 μl fractions, corresponding to 0.9-1.3 ml elution volume, were collected from each sample. The fractions were dried by evaporation in a vacuum centrifuge.

### Liquid chromatography tandem mass spectrometry (LC-MS/MS)

The LC-MS/MS setup consisted of an Easy nLC-1200 HPLC system coupled to an Orbitrap Fusion Lumos mass spectrometer equipped with a Nanospray Flex ion source (all ThermoFisher Scientific). Each SEC fraction was injected in duplicate and samples were separated on an Acclaim PepMap RSLC C18 column (250 mm x 75 μm, 2 Å particle size, ThermoFisher Scientific). Gradient elution was performed using mobile phases A = water/acetonitrile/formic acid (98:2:0.15, v/v/v) and B = acetonitrile/water/formic acid (80:20:0.15, v/v/v) with a gradient of 11 to 40 %B in 60 min and a flow rate of 300 nl/min.

Tandem mass spectra were acquired in the data-dependent acquisition mode with a cycle time of 3 s. Each MS scan was acquired in the orbitrap analyzer at a resolution of 120,000, followed by MS/MS scans in the orbitrap at 30,000 resolution. Precursor ions with a charge state between 3+ and 7+ and an m/z between 350 and 1500 were isolated by quadrupole isolation with an isolation width of 1.2 m/z and fragmented using collision-induced dissociation in the linear ion trap at 35% normalised collision energy. Dynamic exclusion was activated for 30 s after one sequencing event.

### Identification of crosslinked peptides

The identification of crosslinked peptides was performed using xQuest (version 2.1.5, https://gitlab.ethz.ch/leitner_lab/xquest_xprophet)^85^. The data was searched against a database including the proteins of interest plus calmodulin and the most abundant contaminants (a total list of 11 proteins) and a decoy database containing the reversed sequences. The search parameters included trypsin as the enzyme, a maximum of two missed cleavages, carbamidomethylation of Cys as fixed modification, oxidation of Met as variable modification, an MS1 error tolerance of ±15 ppm, and an MS2 error tolerance of ±15 ppm. Subsequently, crosslinked peptide candidates were subjected to a filtering step based on a mass tolerance window of -6 ppm to 1 ppm, a threshold for TIC >0.1, a delta score <0.9, and a minimum of five matched fragment ions per peptide. The resulting datasets correspond to a false discovery rate of less than 1% at the unique peptide-pair level.

### Binding experiment of the NALCN DII-DIII linker peptide and STX1A-SNAP25 complexes

10 μM of synthetic NALCN DII-DIII linker peptide (NALCN amino acids 698-772: Biotin-WGEDNKYIDQKLRKSVFSIRARNLLEKETAVTKILRACTRQRMLSGSFEGQPAKERSILSVQHHIRQERRSLRHGSN-NH2) was incubated with 2.5 μM of either STX1A-SNAP25 wild-type or mutant STX1A-SNAP25-I44R,D51R,E52R,E55R complex for 30 mins on ice in buffer B (25 mM HEPES pH 7.5, 150 mM NaCl and 0.01% (w/v) GDN). The resulting complexes were separated on Superose 6 3.2/300 column and elution fractions were run on SDS-PAGE gels and stained with Coomasie Blue stain.

### Animal maintenance and generation of mouse lines

All procedures for animal maintenance and experiments were in accordance with the regulations of and approved by the animal welfare committee of Charité-Universitätsmedizin and the Berlin state government Agency for Health and Social Services under license number T0220/09. The Stx1b^FL/FL^/Stx1a knockout (KO) mouse was generated by breeding the Stx1a KO line in which exon 2 and 3 are deleted^37^ with the Stx1b conditional KO line in which exon 2–4 are flanked by loxP sites^76^. Infecting neurons with lentiviral *Cre* recombinase leads to complete loss of Stx1a/1b^47^.

### Neuronal cultures

Hippocampal neurons were obtained from mice of either sex at postnatal day (P) 0–2 and seeded on a continental astrocyte feeder layer prepared one to two weeks before neuronal seeding as previously described^47^. Briefly, hippocampi were dissected, and neurons dissociated by an enzymatic treatment using 25 units per ml of papain for 45 min at 37 °C. For electrophysiology and qPCR experiments, hippocampal neurons were seeded at a density of 100 x 10^3^ neurons/well in a 6-well plate, and for survival experiments neurons were plated on an astrocyte feeder layer at a density of 50 x 10^3^ neurons/well in a 12-well plate. The neuronal cultures were incubated for 14–22 (for electrophysiology and qPCR) or 8-22 (for survival analysis) days in-vitro (DIV) in Neurobasal-A supplemented with B-27 (Invitrogen), 50 IU/ml penicillin and 50 µg/ml streptomycin at 37°C before experimental procedures. Neuronal cultures were transduced with lentiviral particles at DIV 1 with between 5×10^5^-1×10^6^ infectious virus units. The viability of the neurons in vitro was defined as the number of surviving neurons at different time points between DIV 8 and DIV 22.

Analysis of neuronal survival was performed as described previously^47^. Phase-contrast brightfield images of 7 randomly selected fields of view (FOV) per well, with 2 wells per group for each culture for a total of 14 FOV per condition and culture, were acquired by an experimenter blinded to the experimental treatment with a DMI4000 microscope, DFC 345 FX camera, HC PL FLUOTAR 20x objectives, and LAS-AF software (all from Leica). Neurons were counted manually with ImageJ software. Transduction with lentiviral constructs was verified by visualising nuclear localization signal (NLS)-green fluorescent protein (GFP) and/or NLS-red fluorescent protein (RFP). To evaluate the rates of cell death, the number of counted neurons at each time point was normalized to the number of neurons counted at DIV 8 for each group. The images presented in Figure 4D have been background-corrected using the ‘rolling ball’ algorithm with a pixel size of 30 and contrast-adjusted using ImageJ^86^ to correct for the uneven illumination as seen in the unprocessed images in Figure S13. The GFP channel uses the OPF fresh lookup table, and the RFP channel uses the OPF orange lookup table, both from https://github.com/cleterrier/ChrisLUTs.

### Design and generation of lentiviral constructs

Lentiviral particles were provided by the Viral Core Facility (VCF) of the Charité-Universitätsmedizin, Berlin, and were prepared as previously described^47^. The cDNA of mouse Stx1a (NM_016801) and the NALCN DII-DIII linker (residue 636 to 861 of XP_011519369) was cloned in frame after an NLS-GFP-P2A (for Stx1a) or a NLS-RFP-P2A (for NALCN) sequence within the FUGW shuttle vector in which the ubiquitin promoter was replaced by the human synapsin 1 promoter (f(syn)w). The improved Cre recombinase (iCre) cDNA was C-terminally fused to RFP-P2A or GFP-P2A for identification of infected cells. To reduce NALCN protein levels in primary hippocampal neurons via a short hairpin (sh)RNA expressing lentivirus, an shRNA cassette containing a 21 bp sense and antisense target sequence of mouse NALCN (5′-GTGCCATCATCAGCGTCATCT-3′) linked by a TCAAGAG linker was cloned downstream of a U6 promoter containing lentiviral shuttle vector that also contained a human synapsin 1 promoter-driven NLS.RFP expression cassette as a reporter (f(U6)shRNA-NALC.hSyn1-NLS.RFP.WPRE).

### Neuronal culture electrophysiology

Whole cell patch-clamp recordings were performed on mass-cultured hippocampal neurons at DIV 14–22 at RT with a Multiclamp 700B amplifier and an Axon Digidata 1550B digitizer controlled by Clampex 10.0 software (both from Molecular Devices). Membrane capacitance and series resistance were compensated by 70% and only the recordings with a series resistance smaller than 10 MΩ were used for further experiments. Data were sampled at 10 kHz and filtered with a low-pass Bessel filter at 3 kHz. Extracellular solution was constantly perfused and contained the following: 140 mM NaCl, 3 mM KCl, 10 mM HEPES, 10 mM glucose, 2 mM CaCl_2_ and 2 mM MgCl_2_ (adjusted to 300 mOsm with D-glucose; pH 7.4 with NaOH). Borosilicate glass patch pipettes were pulled with a multistep puller (Sutter Instruments), yielding a final tip resistance of 2 – 5 MΩ when filled with intracellular solution containing the following: 104 mM CsCH_3_SO_3_, 30 mM TEACl, 10 mM HEPES, 10 mM EGTA, 1 mM MgCl_2_, 3 mM Na_2_ATP, 0.3 mM Na_2_GTP (290 mOsm; pH 7.4 with CsOH). To monitor Ca^2+^-sensitive sodium leak currents, neurons were clamped at –70 mV and holding currents were recorded as a fast perfusion system (SF-77B; Warner Instruments) was deployed to rapidly switch the extracellular solution around the patched neuron. The solutions used were either the standard extracellular solution as described above, with lowered divalents (0.5 mM CaCl_2_ and 0.5 mM MgCl_2_), or with lowered divalents and with NMDG^+^ instead of Na^+^ (140 NMDG^+^, 0.5 mM CaCl_2_ and 0.5 mM MgCl_2_). Each of these three solutions were supplemented with 30 µM strychnine, 10 µM bicuculline, 10 µM NBQX, and 1 µM TTX to block ligand-gated synaptic channels and voltage-gated sodium channels.

### Statistical analysis and visualisation

Statistical models were fit in R using the brms package^87^ and visualised with the tidybayes package^88^, and unless otherwise stated were fit with multilevel Bayesian regression models. The Boltzmann equation 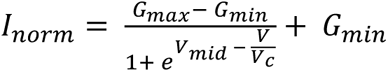 was used to model the conductance-voltage relationships in Figure 2D and Figure S4B, with *V*_*mid*_ and *V*_*C*_ estimated as group level effects which varied between oocytes. The same equation was used to fit the conductance-voltage relationships for the Na_V_ recordings in Figure S8A, but with *G*_*min*_and *G*_*max*_fixed to 0 and 1 respectively. Similarly, the current deactivation traces in the same figures were fit to the equation *I*_*norm*_ = (*I*_*initial*_ − *I*_*final*_) · *e*^1/*tau* ·*s*^ + *I*_*final*_, with *I*_*final*_ and *tau* estimated as group level effects which varied between oocytes. The fit summaries are presented as the overall (or population level) median estimate and the 95% quantiles, with the group level individual oocyte parameters shown as data points in Figure S4B.

The calcium sensitive sodium leak data from neurons in Figure 4B were fit to a linear regression model with a population level intercept for each condition, and the residual standard deviation allowed to vary between conditions (as the variance of the data was not equal between conditions). Contrasts between the population level estimates for each condition were calculated as the pairwise differences between the intercept distributions, and the full distributions of the contrasts are shown together with the median, 66%, and 95% quantiles. Similarly, the proportion of surviving neurons at DIV22 were fit to a linear regression model with a population level intercept for each condition. The full distributions of the resulting fits are shown together with the median, 66%, and 95% quantiles.

## Acknowledgements

We thank members of the Pless and Rosenmund labs for discussion and advice during the preparation of this manuscript, Paola Picotti (ETH Zurich) for access to instrumentation and laboratory infrastructure, Genentech colleagues in the BioMolecular Resources and Structural Biology departments for their support, and appreciate the encouragement of C. Ciferri, M. Masureel, V. Dixit, A. Chan and K. Briner. We would also like to thank Dr. Han Chow Chua and Dr. Cordelia Imig for their critical reading of the manuscript.

## Author contributions

Conceptualization, S.U., E.T., A.S.H., A.L., C.R., M.K., S.P.; Formal Analysis, S.U., R.F., S.Y., C.C.J, M.K.; Investigation, S.U., E.T., R.F., S.Y., C.C.J.; Resources, J.C., P.J., I.Z., T.T., B.B., Writing – Original Draft, S.U.; Writing – Review and Editing, E.T., R.F., A.S.H, A.L., C.R., M.K., S.P.; Visualization, S.U., E.T., R.F., S.Y., C.C.J., A.L., M.K.; Supervision, E.T., A.S.H., A.L., C.R., M.K., S.P.; Project Administration, A.L., C.R., M.K., S.P.

## Competing Interest Statement

S.Y., C.C.J., P.J., I.Z. and M.K. are employees of Genentech/Roche. The other authors declare to have no competing interests.

## Data and code availability

All data needed to evaluate the conclusions in the paper are present in the paper, the Supplementary Materials and the below repositories. Code to reproduce the computational screening steps can be found at https://github.com/smusher/NALCN_protein_screen. The mass spectrometry proteomics data have been deposited to the ProteomeXchange Consortium via the PRIDE^89^ partner repository with the dataset identifier PXD053999. All other raw data and associated analysis scripts have been uploaded to Zenodo with the DOI: 10.5281/zenodo.13149886

## Funding

This work was supported by the Carlsberg Foundation (grant CF20-0248), the Independent Research Fund Denmark (1026-00325A and 7025-00097A) and the Lundbeck Foundation (R383-2022-165).

## Notes

### Summary of Updates

Added missing author to bioRxiv submission. The author was always correctly included in the manuscript PDF.

