## Supplemental Information for "The sodium leak channel NALCN is regulated by neuronal SNARE complex proteins"

### Supplementary figures S1-S13

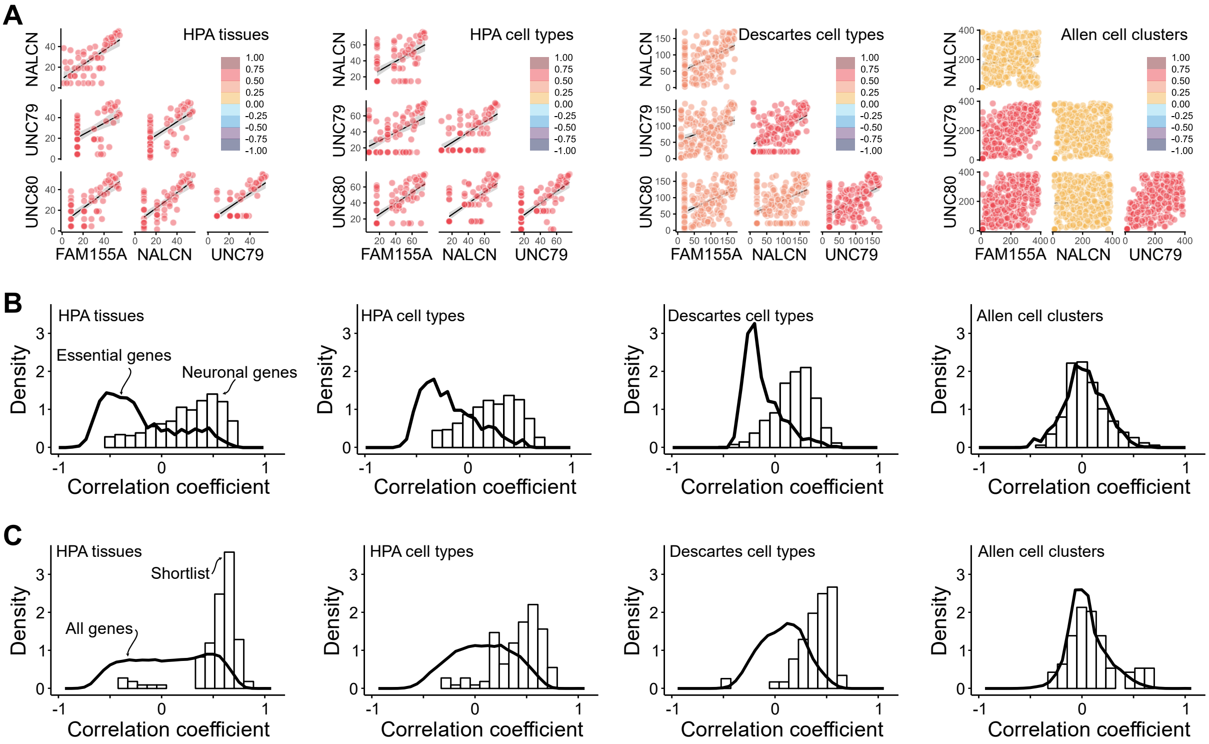


**Figure S1. Transcript expression correlation and frequency distributions across the 4 datasets used.** A) Transcript expression correlation between the NALCN core complex genes across the 4 datasets used. Points are colour-coded by the pairwise Spearman's rank correlation coefficient. B) Frequency distributions of correlation coefficients for essential genes (lines) and genes linked to neuronal function (histograms). C) Frequency distributions of correlation coefficients for all protein-coding genes (lines) and the 25 gene functional screening list (histograms).

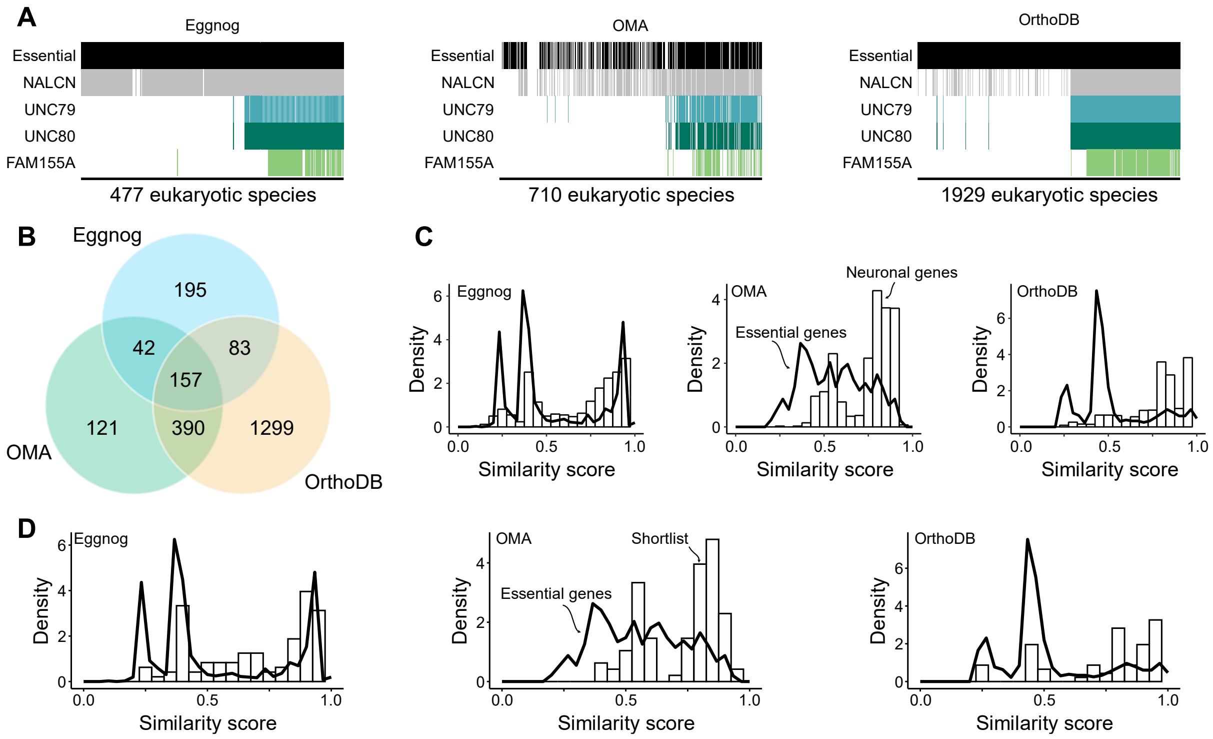


**Figure S2. Phylogenetic fingerprints, overlap and frequency distributions for the 3 databases used.** A) Phylogenetic fingerprints for the NALCN core complex genes and the modal fingerprint for essential genes for the 3 databases used. B) Overlap of eukaryotic species present in each of the 3 databases. C, D) Frequency distributions of phylogenetic fingerprint similarity scores between essential genes (lines) and either genes linked to neuronal function (histograms panel C) or the 25 gene functional screening list (histograms panel D).

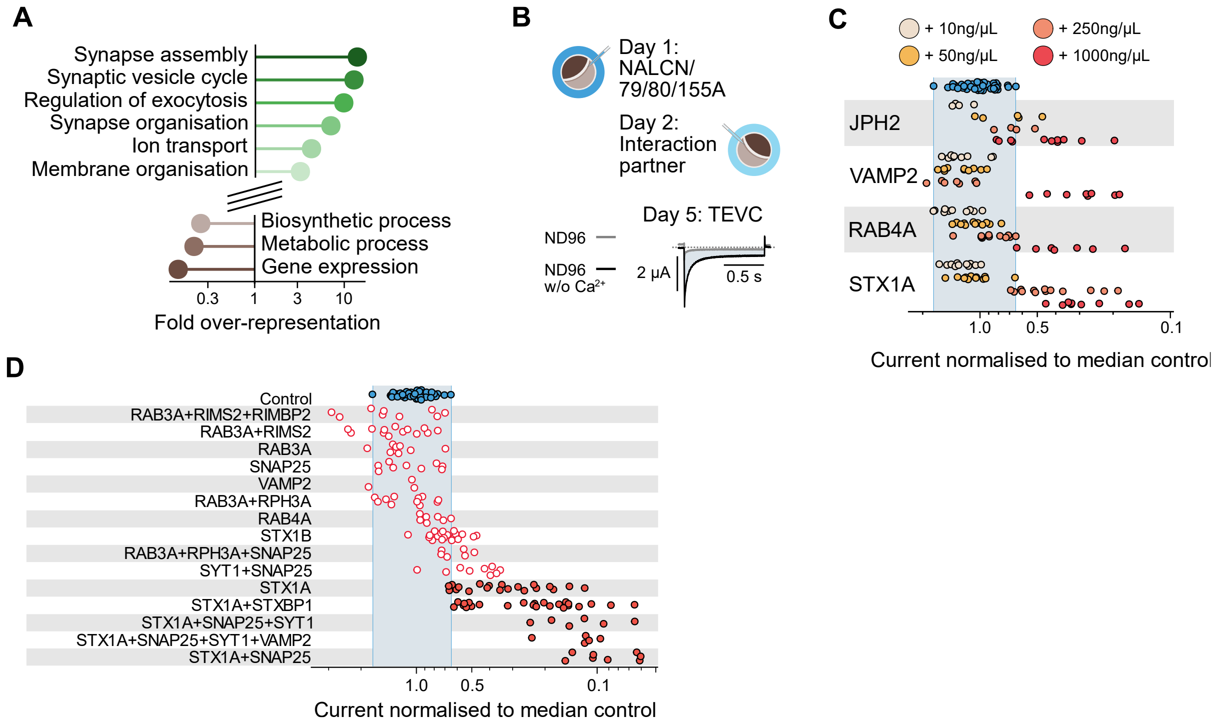


**Figure S3. Overrepresented GO-terms and functional screening of shortlist.** A) Over-representation of GO-terms in the 200 genes most similar to the NALCN core complex (inside circle of Figure 1C) against background (outside of circle). The six most over-represented and the three most under-represented terms are shown here. B) Overview of the shortlist functional screening process. C) Oocytes were injected, and data were analysed as in Figure 1E, but with a range of mRNA concentrations for the four candidate proteins. Data points from injection with 1000 ng/µL are reproduced from Figure 1E. D) Combinations of protein-encoding mRNAs were injected together and data were analysed as in the previous panel.

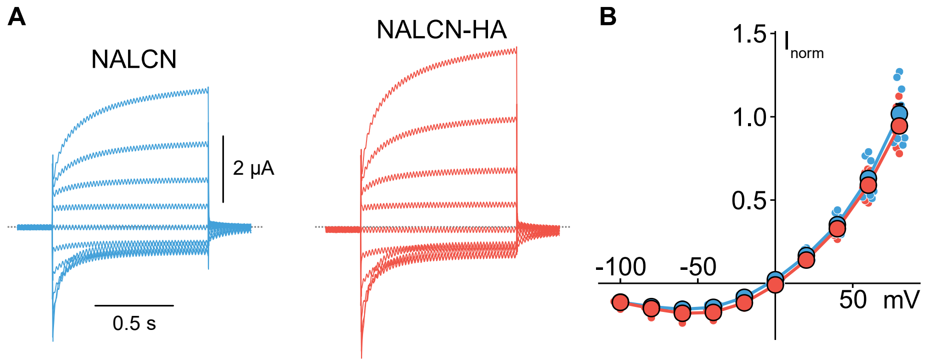


**Figure S4. Functional characterization of wild-type and HA-tagged NALCN.** A) Example traces from wild-type NALCN (left, blue) or HA-tagged NALCN (right, red) both expressed with UNC79-UNC80-FAM155A. B) Steady-state current-voltage relationships from the same conditions as in panel A.

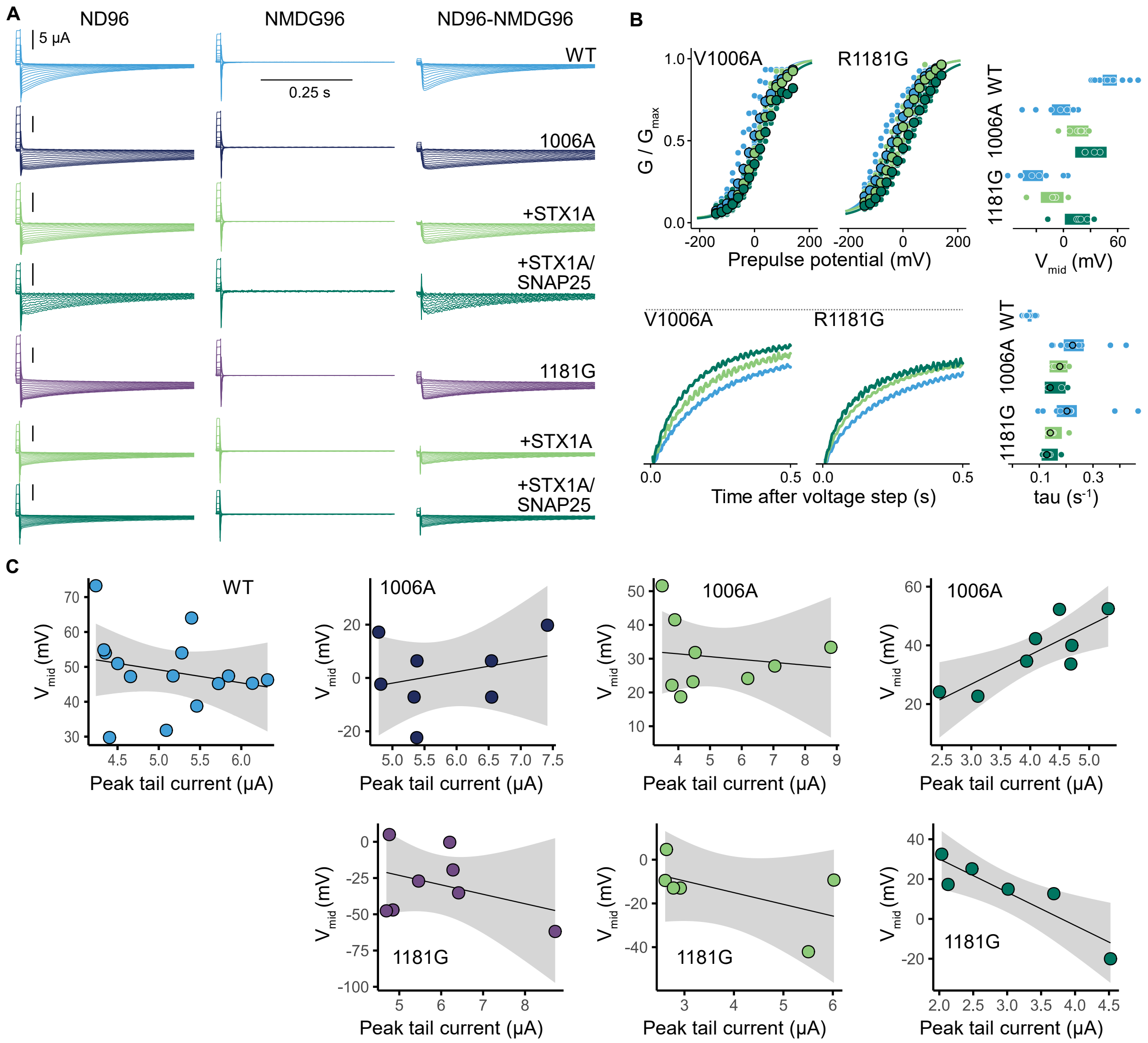


**Figure S5. Tail current recordings.** A) Example traces from tail current recordings used in Figure 2E and in panel B of this figure. Left: first recording in ND96 w/o $Ca^{2+}$, middle: second recording in NMDG96, right: subtracted traces. B) Upper: conductance-voltage curves from *Xenopus* oocytes expressing the indicated gain-of-function NALCN point mutants expressed either with UNC79-UNC80-FAM155A alone (blue), or additionally expressing STX1A (light green) or STX1A and SNAP25 (dark green). The $V_{mid}$ estimates for each oocyte are shown as data points with white outlines in the furthest right panel, with the 95% credible intervals of the group-level estimates shown as coloured blocks. Lower: representative current deactivation traces for the indicated gain-of-function NALCN point mutants expressed either with UNC79-UNC80-FAM155A alone (blue), or additionally expressing STX1A (light green) or STX1A and SNAP25 (dark green). Grey dashed lines indicate the zero current level. Right: tau estimates from single exponential fits to traces from individual oocytes are shown as data points with white outlines, with the 95% credible intervals of the group-level estimates shown as coloured blocks. The tau values for the displayed traces are outlined in black. C) Relationship between current magnitude and estimated $V_{mid}$. Each panel shows data from a different construct and coexpression, with colours the same as in panel A. The black line and grey area are the point estimate and standard error from fits to a linear regression.

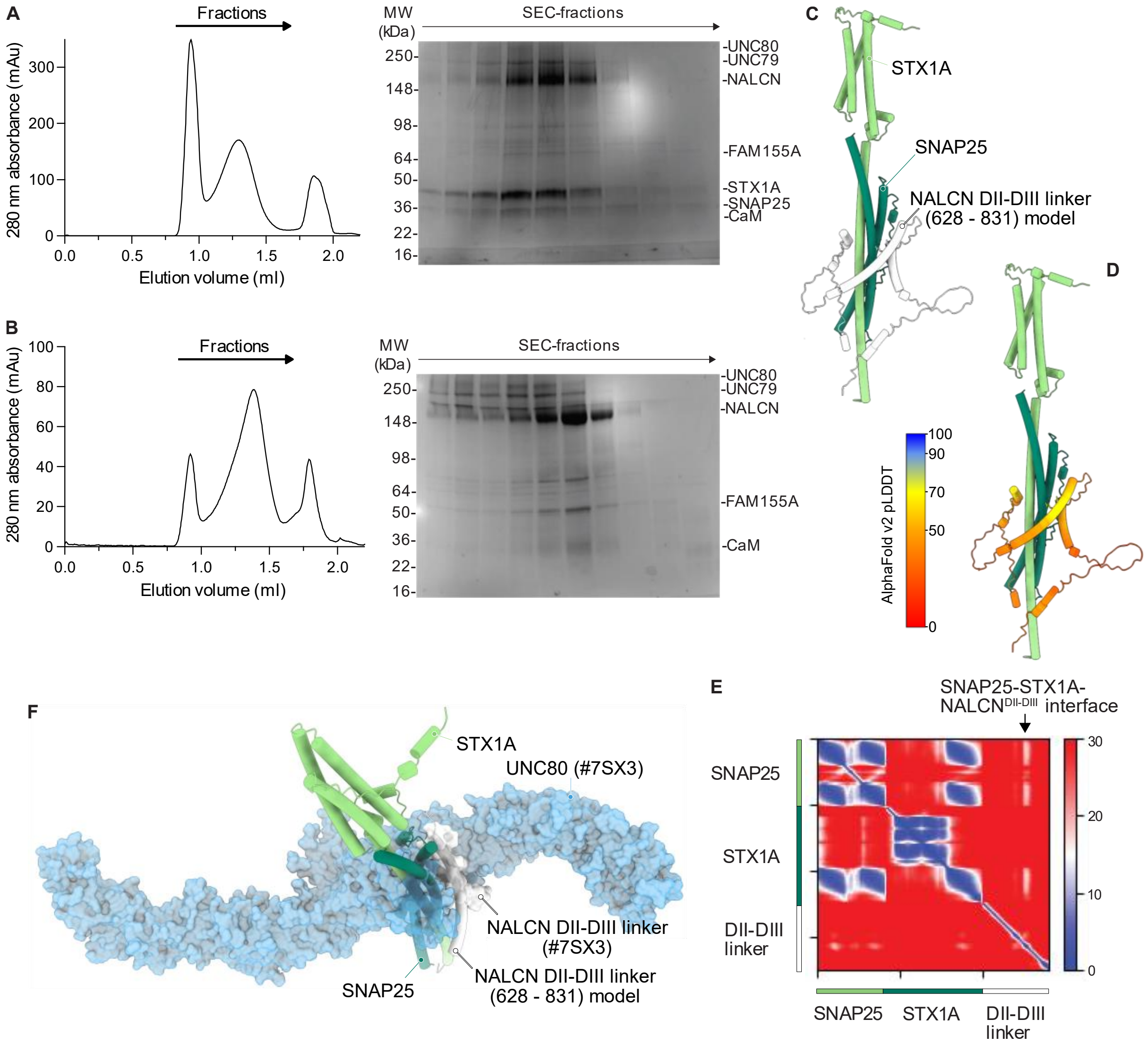


**Figure S6. Biochemical and computational analysis of the NALCN-STX1A-SNAP25 interaction.** A) Size exclusion chromatography and corresponding SDS-PAGE of the purified NALCN-FAM155A-STX1A-SNAP25 complex co-expressed with UNC79 and UNC80 and B) of the NALCN-FAM155A-UNC79-UNC80 complex. C) AlphaFold v2 multimer prediction of STX1A-SNAP25 and the NALCN-DII-DIII linker (P628-P831) region. D) Per-residue model confidence score (pLDDT) mapped on the model prediction described in B. dark blue: pLDDT > 90, light blue: pLDDT between 90-70, yellow: pLDDT between 70-50, red: pLDDT <50 E) Predicted aligned error (PAE) of the model described in B. F) Overlay of UNC80-NALCN-DII-DIII linker from the NALCN-FAM155A-UNC80-UNC79-CaM co-structure (PDB:7SX3) and predicted STX1A-SNAP25-NALCN-DII-DIII linker. Models aligned on NALCN DII-DIII linker region V711-S740.

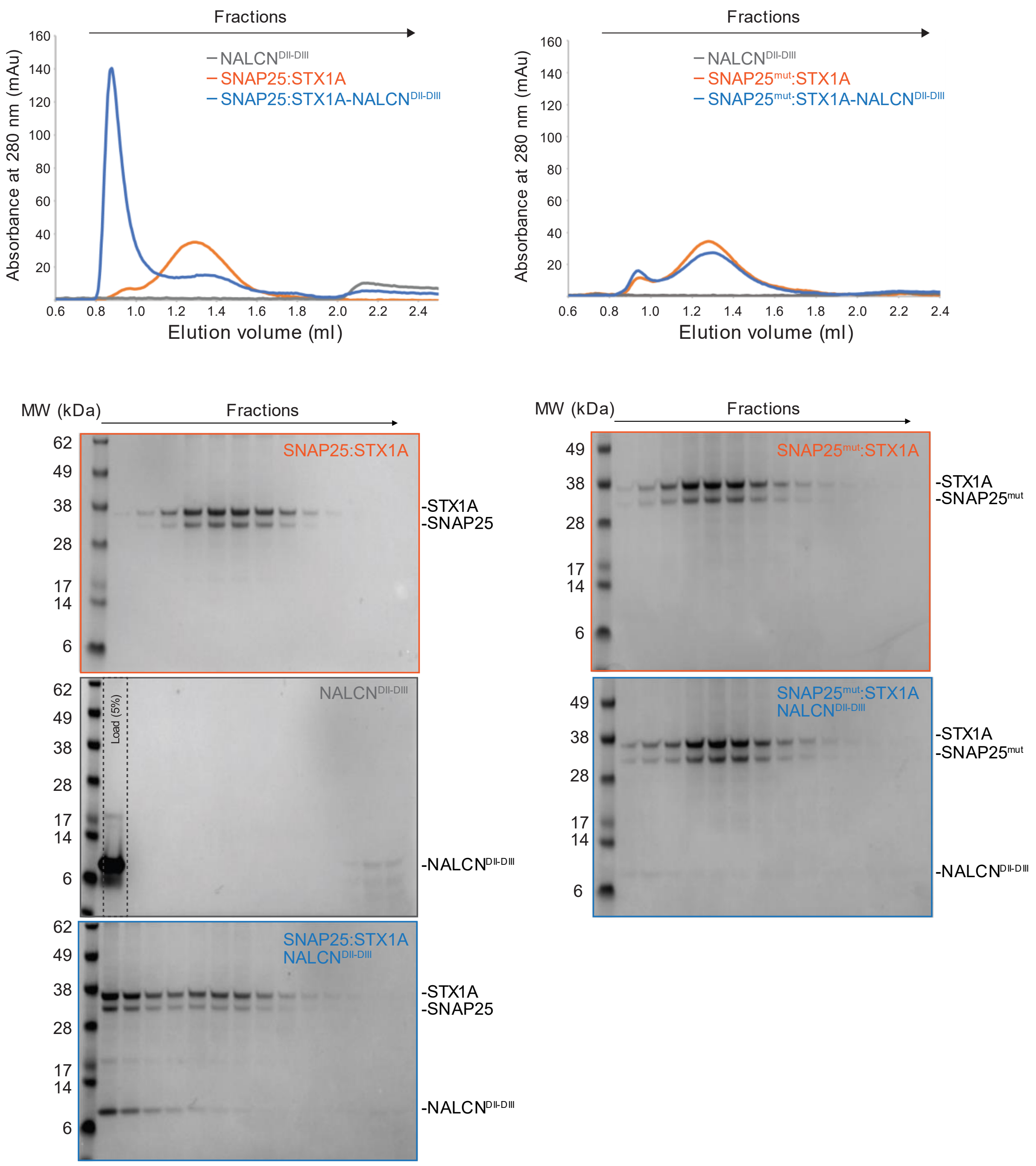


**Figure S7. Biochemical characterisation of the NALCN-STX1A-SNAP25 interaction.** Size exclusion chromatography (top) and corresponding SDS-PAGE (bottom) of individual and complexed NALCN DII-DIII linker (E698-N772) peptide with STX1A-SNAP25 (left) or with STX1A-SNAP25-44/51/52/55R (SNAP25$^{mut}$; right).

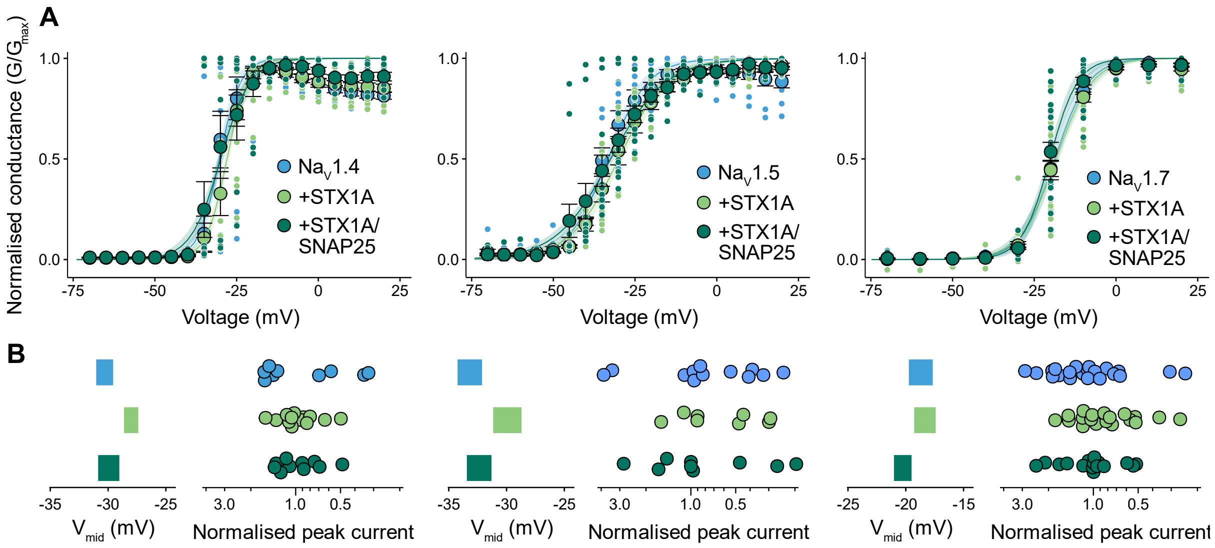


**Figure S8. Conductance-voltage relationships of $Na_V1.4$, $Na_V1.5$ and $Na_V1.7$ with STX1A +/- SNAP25.** A) Conductance-voltage relationships from three different voltage-gated sodium channels expressed alone or in combination with STX1A +/- SNAP25. Fitted curves are to a Boltzmann equation. B) Left: 95% credible intervals of the group-level estimates for the $V_{mid}$ parameters of the curves in A, coloured to match. Right: Summary peak current magnitudes from the same recordings normalised to the median of the $Na_V$ alone from the same oocyte batch.

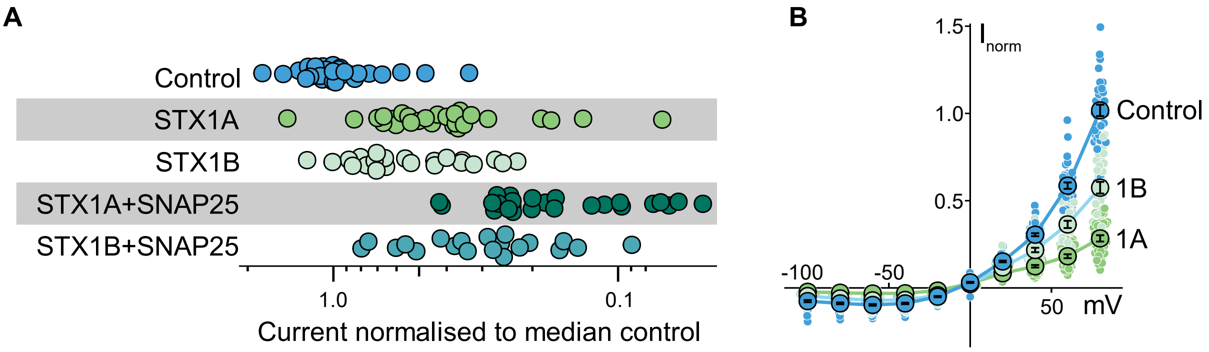


**Figure S9. STX1A/B have similar functional effects.** A) Summary current magnitudes from cells expressing NALCN-UNC79-UNC80-FAM155A alone or in combination with STX1A/B +/- SNAP25. B) Steady-state current-voltage relationships from the same conditions as in panel A, with points sharing the same colour scheme.

**A**

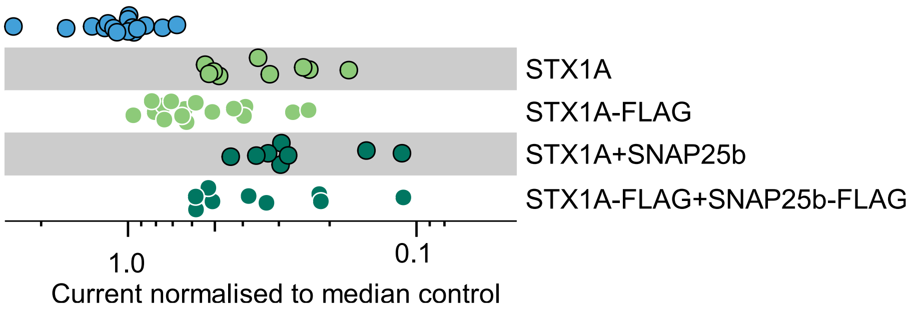


**Figure S10. Impact of FLAG-tag on STX1A inhibition of NALCN currents.** Summary current magnitudes from *Xenopus* oocytes expressing NALCN/UNC79/UNC80/FAM155A alone (blue), and combinations of wild-type or FLAG-tagged STX1A and SNAP25.

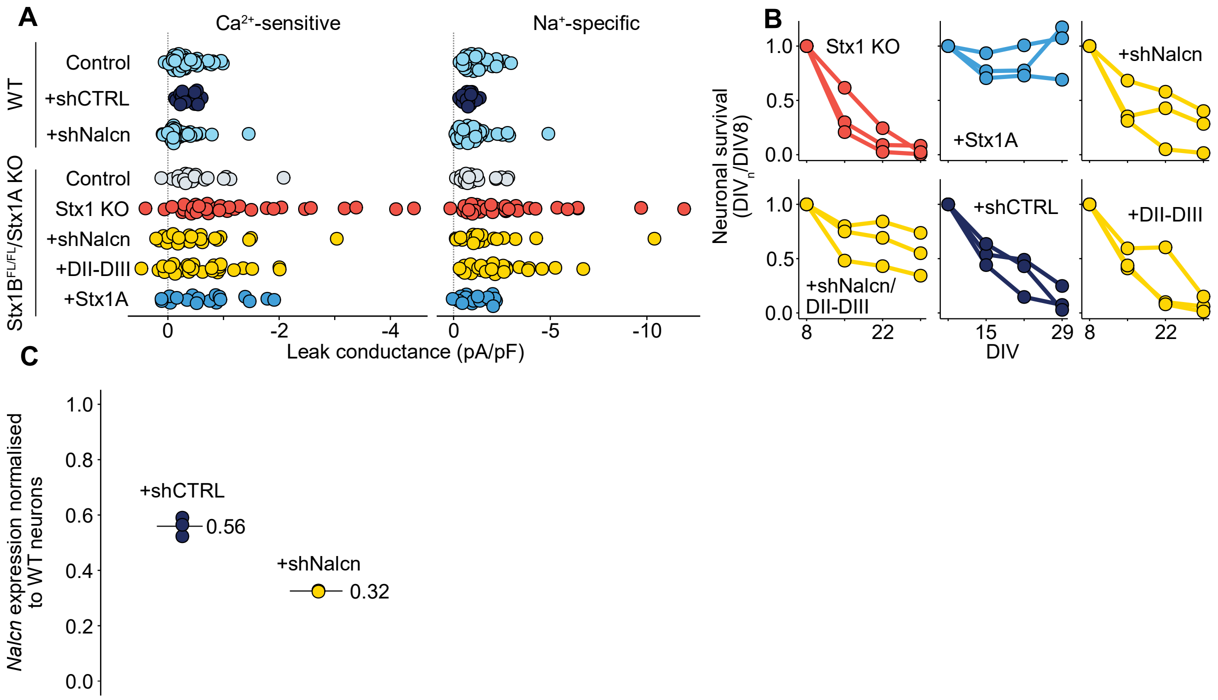


**Figure S11. Leak current and neuronal survival summaries.** A) $Ca^{2+}$-sensitive and $Na^{+}$-specific composition of neuronal leak currents. Data in the left panel are the same as Figure 4A but are shown here on a normal (not log) x-axis. Data in the right panel are calculated by subtracting the magnitude of the holding current in $NMDG^{+}$ based extracellular solution from that in $Na^{+}$ based extracellular solution. B) Full results from the experiment shown in Figure 4C. Each panel contains the results of three separate neuronal cultures infected with the lentiviral constructs indicated. C) shNalcn efficacy assayed by RT-qPCR from hippocampal neurons as described in the methods. Each point is a biological replicate from a separate culture.

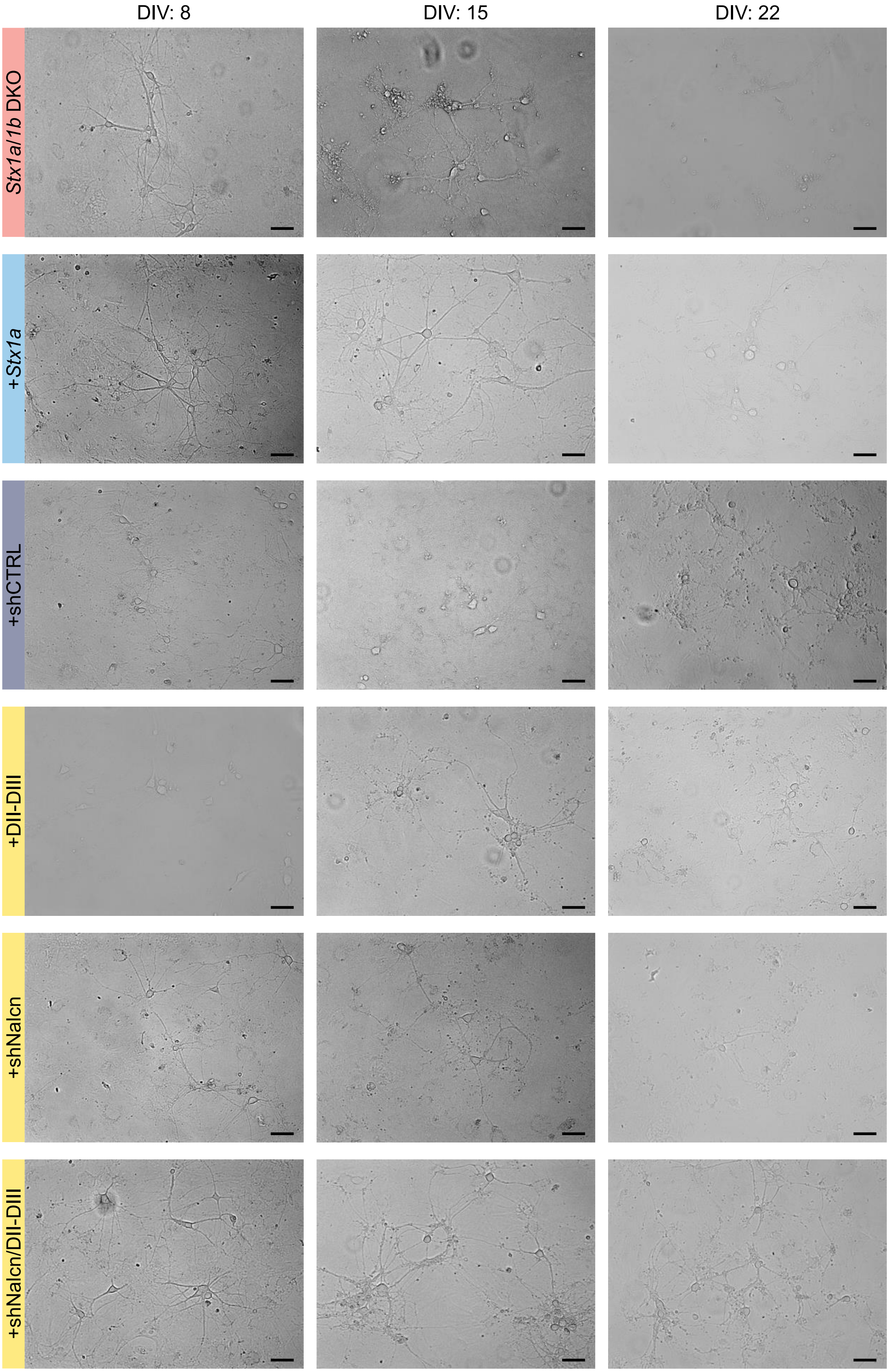
DIV: 8
DIV: 15
DIV: 22
Stx1a/1b DKO
+Stx1a
+shCTRL
+DII-DIII
+shNalcn
+shNalcn/DII-DIII

**Figure S12. Representative brightfield images for neuronal survival experiment shown in Figure S11B.** Representative brightfield images for each imaging stage and each culture from the neuronal survival experiment shown in Figure S11B. These images were not altered after acquisition. Scale bars are 50 μm.

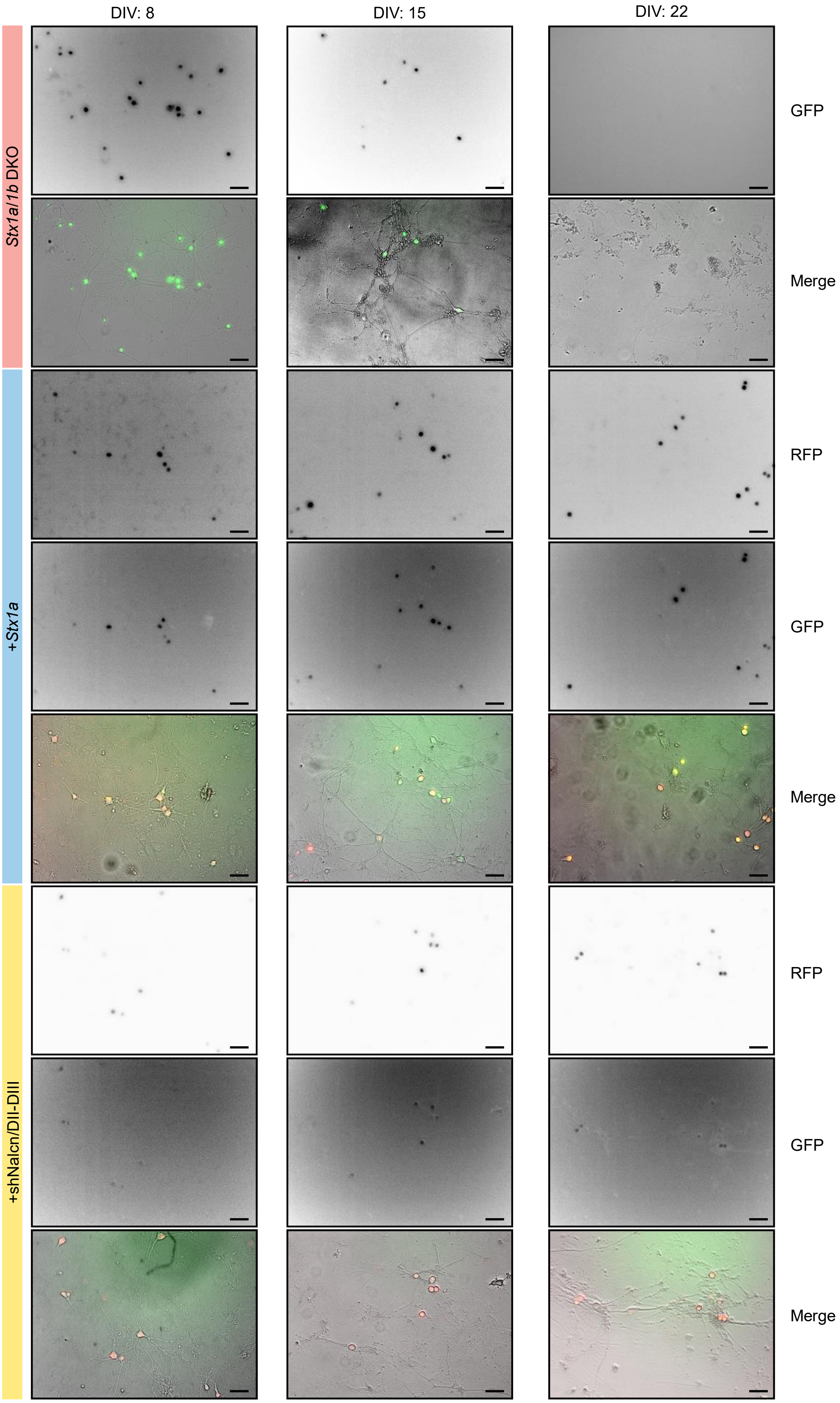
DIV: 8
DIV: 15
DIV: 22
Stx1a/1b DKO
+Stx1a
+shNalcn/DII-DIII
GFP
Merge
RFP
GFP
Merge
RFP
GFP
Merge

**Figure S13. Representative brightfield images and fluorescent images for neuronal survival experiment shown in Figure 4D.** Representative brightfield and fluorescent images from each imaging stage and each culture from the neuronal survival experiment shown in Figure 4D. Fluorescent images are shown with an inverted lookup table. Scale bars are 50 μm.
